# Ovarian disrupting effects and mechanisms of long- and short-chain per- and polyfluoroalkyl substances in mice

**DOI:** 10.1101/2024.02.20.581034

**Authors:** Pawat Pattarawat, Tingjie Zhan, Yihan Fan, Jiyang Zhang, Hilly Yang, Ying Zhang, Sarahna Moyd, Nataki C. Douglas, Margrit Urbanek, Brian Buckley, Joanna Burdette, Qiang Zhang, Ji-Yong Julie Kim, Shuo Xiao

**Affiliations:** Department of Pharmacology and Toxicology, Ernest Mario School of Pharmacy, Rutgers University, Piscataway, NJ 08854, USA; Environmental and Occupational Health Sciences Institute (EOHSI), Rutgers University, Piscataway, NJ 08854, USA; Center for Environmental Exposures and Disease, Rutgers University, Piscataway, NJ 08854, USA; Department of Epidemiology, Rollins School of Public Health, Emory University, Atlanta GA 30322, USA; Gangarosa Department of Environmental Health, Rollins School of Public Health, Emory University, Atlanta GA 30322, USA; Department of Obstetrics, Gynecology and Reproductive Health, New Jersey Medical School (NJMS), Rutgers University, Newark, NJ 07103; Center for Immunity and Inflammation, Rutgers Biomedical and Health Sciences (RBHS), Newark, NJ 07103, USA; Department of Medicine, Northwestern University Feinberg School of Medicine, Chicago, IL, USA; Department of Pharmaceutical Biosciences, Center for Biomolecular Science, University of Illinois at Chicago, Chicago, IL 60607, USA; Department of Obstetrics & Gynecology, Feinberg School of Medicine, Northwestern University, Chicago, IL 60611, USA

**Keywords:** Per- and polyfluoroalkyl substances, long-chain PFAS, short-chain PFAS, alternative, follicle maturation, ovulation, ovarian hormone secretion, peroxisome proliferator-activated receptor gamma

## Abstract

**Background:** The extensive use of per- and polyfluoroalkyl substances (PFAS) has led to environmental contamination and bioaccumulation. Previous research linked PFAS exposure to female reproductive disorders, but the mechanism remains elusive. Further, most studies focused on legacy long-chain PFOA and PFOS, yet the reproductive impacts of other long-chain PFAS and short-chain alternatives are rarely explored.

**Objectives:** We investigated the effects and mechanisms of long- and short-chain PFAS on the ovary and associated ovarian functions.

**Methods:** A 3D *in vitro* ovarian follicle culture system and an *in vivo* mouse model, together with approaches of reverse transcription-quantitative polymerase chain reaction, enzyme-linked immunosorbent assay, RNA-sequencing, pharmacological treatment, *in situ* zymography, histology, *in situ* hybridization, analytical chemistry, and benchmark dose modeling (BMD), were used to test environmentally relevant exposure levels of six long- and short-chain PFAS on follicle maturation, hormone secretion, and ovulation.

**Results:** *In vitro* exposure revealed that long-but not short-chain PFAS interfered with gonadotropin-dependent follicle maturation, ovulation, and hormone secretion. Mechanistically, long-chain perfluorononanoic acid (PFNA) acted as a peroxisome proliferator-activated receptor gamma (PPAR*γ*) agonist in granulosa cells to disrupt follicle-stimulating hormone (FSH)-dependent follicle maturation, luteinizing hormone (LH)-stimulated ovulation, and associated gene regulatory pathways. *In vivo* mouse exposure confirmed the ovarian accumulation of PFNA and the mechanism of PPAR*γ*-mediated ovarian toxicities of PFNA observed *in vitro*. The BMD analysis of *in vitro* and *in vivo* results suggested human relevant exposure levels of long-chain PFAS in our study pose an extra risk of ovarian defects, with follicular rupture as the most sensitive endpoint.

**Discussion:** Using *in vitro* follicle culture and *in vivo* mouse models, we discovered that long-chain PFAS interfere with gonadotropin-dependent follicle maturation, hormone secretion, and ovulation, posing a non-negligible risk to women’s reproductive health including anovulation, irregular menstrual cycles, and sub- or infertility.

## Introduction

The per– and polyfluoroalkyl substances (PFAS) are thousands of synthetic compounds that consist of a major carbon backbone and at least one fluoroalkyl moiety (C_n_F_2n+1_-) [1]. PFAS that contain 8 or more carbons are defined as long-chain PFAS, including perfluorooctanoic acid (PFOA), perfluorooctane sulfonate (PFOS), and perfluorononanoic acid (PFNA). Short-chain PFAS contain 4-7 carbons, such as perfluorobutane sulfonic acid (PFBS), perfluoroheptanoic acid (PFHpA), and GenX, and are now increasingly manufactured and applied as alternatives [2, 3]. The strong C-F bonds of PFAS render them with low surface tension and thermally and oxidatively resistant [4]. Since 1960s, PFAS have been widely used in numerous consumer and industrial products, including textiles, food packaging, cookware coating, surfactants, personal care products, and aqueous firefighting foams [5–7]. PFAS are highly resistant to environmental degradation, which has led to extensive environmental contamination and bioaccumulation [8, 9] and earn the name “Forever Chemicals” [4, 10]. Due to their health threats to human and wildlife animals, there is a growing global environmental and public health concern [11].

Humans are exposed to PFAS primarily through drinking water, while it is also possible through contaminated foods, soil, contact with PFAS-containing products, and PFAS manufacturing process [12–16]. These exposures impact millions of residents in the US and the larger population worldwide [17–19]. The strong C-F bonds make many PFAS rarely metabolized in human bodies after absorption, particularly the long-chain PFAS, leading to prolonged half-lives up to 8-9 years [20, 21], while short-chain PFAS in general have shorter half-lives [22]. Long-chain PFAS are detectable in > 90% of world populations [23–25], with blood concentrations ranging from 1.5 to 220 µM [20, 23, 25–30].

Exposure to PFAS has been related to several adverse health outcomes in humans, such as the induction or promotion of liver toxicity [31, 32], cancers [33], immunotoxicity [34, 35], and metabolic disorders [36]. While the adverse impacts of PFAS on reproductive health remains controversial [37–39], growing epidemiological evidence suggests associations between PFAS exposure and female reproductive dysfunctions related to the ovary. These include premature ovarian failure (POF) [40, 41], irregular menstrual cycles [42], polycystic ovary syndrome (PCOS) [43, 44], and infertility [45–47]. The ovary houses follicles at various stages to sustain female reproductive cycles, fertility, and overall health. The early phase of follicle activation and development is largely gonadotropin independent, but the eventual maturation of a secondary or early antral follicle is governed by the pituitary gonadotropin follicle-stimulating hormone (FSH), and the ovulation of a fully mature preovulatory follicle is triggered by another pituitary gonadotropin, luteinizing hormone (LH) [48–51]. Experimental studies reported that oral exposure to long-chain PFOA and PFOS in rodents at 1 –10 mg/kg (0.045 – 0.45 mg/kg human equivalent dose) adversely impacted ovarian cyclicity [52–55], steroidogenesis [56–58], and oocyte maturation [59–62]. However, the underlying mechanism remains elusive. Moreover, nearly all previous studies focused on PFOA and PFOS, the two long-chain legacy PFAS that have been gradually phased out in the US and many other counties. Other long-chain PFAS (e.g., PFNA) and emerging short-chain PFAS (e.g., GenX and PFBS), however, may reach similar contamination levels [2, 3, 63], but few examined their impacts on female ovarian functions and reproduction.

The objective of this study was to investigate the ovarian disrupting effects of PFAS and the toxic mechanisms involved. Long-chain PFAS have been shown to exhibit longer half-lives after absorption and stronger binding affinities for target proteins, such as the peroxisome proliferator-activated receptors (PPARs), than short-chain PFAS [22, 64, 65]. PFAS, particularly long-chain PFAS, have been speculated to exhibit endocrine-disrupting effects by activating PPARs [65, 66]. There are three PPAR subtypes: PPARα, PPARβ/δ (referred to as PPAR*β* below), and PPAR*γ*. PPAR*γ* is primarily expressed in mural granulosa cells of maturing and preovulatory follicles and has been shown to regulate gonadotropin-dependent follicle maturation and ovulation [67–72]. These facts motivated us to hypothesize that PFAS act as a PPAR*γ* agonist in follicular granulosa cells, as the molecular initiating event (MIE), to interfere with gonadotropin-dependent follicular functions, and exposure to long– and short-chain PFAS may exhibit differential ovarian disrupting effects. To test this hypothesis, we first used a 3D *in vitro* mouse ovarian follicle culture system to test for various human relevant concentrations of six long– and short-chain PFAS on gonadotropin-dependent follicle maturation, hormone secretion, and ovulation, and determined the ovarian toxic effects and dose-response relationship of the exposed PFAS. PFNA was then selected for *in vitro* exposures to determine the molecular mechanisms involved and for *in vivo* exposures in mice to verify its accumulation in the ovary and ovarian toxic effects identified *in vitro*. Both *in vitro* and *in vivo* exposure models demonstrated that long-chain PFAS are more ovarian toxic than short-chain PFAS, and PFNA acts as an PPAR*γ* agonist in follicular granulosa cells to interfere with gonadotropin-dependent follicle maturation, hormone secretion, and ovulation, posing a non-negligible risk to women’s reproductive health by heightening the probability of anovulation, irregular menstrual cycles, and sub– or infertility.

## Materials and Methods

### Animals

Both CD-1 mouse breeding colony for ovarian follicle isolation and *in vitro* PFAS exposure and young adult CD-1 female mice (Envigo, Indianapolis, IN) for *in vivo* PFAS exposure were housed in polypropylene cages in the Animal Care Facility of Research Tower at Rutgers University. Mice were kept under a temperature–, humidity–, and light– (12/12 light/dark cycle) controlled facilities and were provided with food and water ad libitum. All animals were maintained and treated according to the NIH Guideline for the Care and Use of Laboratory Animals and the approved Institutional Animal Care and Use Committee (IACUC) protocol at the Rutgers University [73].

### PFAS

PFOA, PFOS, PFNA, PFHpA, and PFBS were purchased from Sigma –Aldrich (St. Louis, MO). GenX was purchased from Manchester organics (Cheshire, UK). Stock solution for PFOA, PFOS, PFNA, PFHpA and PFBS were prepared in DMSO. Stock solution for GenX was prepared in UltraPure water because it has been reported to be degradable in DMSO [74–76]. PFAS were diluted in follicle culture media for *in vitro* exposure and in PBS for *in vivo* exposure, with the concentration of DMSO at 0.1%.

### *In vitro* 3D encapsulated follicle growth (eIVFG), ovulation, and PFAS exposure

Mouse ovaries were collected from 16-day-old CD-1 female mice and then incubated in the enzymatic digesting solution containing L15 media (Invitrogen, Carlsbad, CA) with 30.8 µg/mL liberase and 200 µg/mL DNase for 25 minutes. Immature preantral follicles of the size of 150-180 µM were isolated. This procedure enabled us to obtain ∼150 follicles from 3-5 prepubertal mice. Isolated follicles were encapsulated for *in vitro* culture as we previously described [77–81]. Briefly, follicles were encapsulated in 0.5% (w/v) alginate hydrogel (Sigma-Aldrich, St. Louis, MO). Encapsulated follicles were individually cultured in 96-well plates with each well containing a single follicle and 100 μL growth media consisting of 50% αMEM Glutamax (Thermo Fisher Scientific) and 50% F-12 Glutamax (Thermo Fisher Scientific) supplemented with 3 mg/ml bovine serum albumin (BSA; Sigma-Aldrich), 10 mIU/mL human recombinant follicle stimulating hormone (rFSH; gifted from Dr. Mary Zelinski from the Oregon Non-human Primate Research Center at the Oregon Health and Science University, Beaverton, OR, USA), 1 mg/mL bovine fetuin (Sigma-Aldrich), and 5 μg/mL insulin-transferrin-selenium (ITS, Sigma-Aldrich). Follicles were cultured for 6 days at 37 °C in a humidified environment of 5% CO_2_ in air, which allowed immature follicles to grow from the multi-layered secondary stage to the antral stage to reach maturation.

On day 6 of eIVFG, grown mature antral follicles were freed from alginate through incubation in L15 media consisting of 1% FBS and 10 IU/mL alginate lyase (Sigma-Aldrich) at 37 °C for 10 minutes. Follicles were then incubated in the ovulation induction media containing 50% αMEM Glutamax and 50% F-12 Glutamax supplemented with 10%FBS, and 1.5 IU/mL hCG (Sigma-Aldrich), and 10 mIU/mL rFSH. Ovulation was assessed at 14 hours post-hCG. A follicle was defined as “ruptured” when one side of the follicular wall was breached and defined as “unruptured” when the follicular wall was intact. Oocytes released from ruptured follicles were examined for the extrusion of the first polar body, and oocytes with the first polar body extrusion were defined as MII oocytes. For those non-MII oocytes, most of them were at the germinal vesicle breakdown (GVBD) stage, and a few were at the germinal vesicle (GV) stage. We defined those immature oocytes at GVBD or GV stage as non-MII oocytes. Follicles treated with hCG for ovulation induction were continuously cultured in the ovulation induction media without FSH for 48 hours to induce luteinization and progesterone secretion. The conditioned media was collected at 48 hours and stored at –20 °C for subsequent measuring of progesterone using ELISA.

Cultured follicles were randomly allocated into different experimental groups with 10-15 follicles in each group. In Tier 1 ovarian toxicity testing, follicles were exposed to 1, 10, 100, and 250 μM of long-chain PFAS (PFOA, PFOS, or PFNA) or short-chain PFAS (PFHpA, PFBS, or GenX) during both the follicle maturation window (day 2-6) and the ovulation window (day 6-7). In Tier 2 or 3, follicles were either exposed to PFAS during the follicle maturation window or the ovulation window (day 6-7). The concentration range of PFAS was determined based on the fact that human blood concentrations of PFAS, particularly long-chain PFAS, range from 0.007 – 92.3 µg/mL or 16.9 nM – 222 µM, depending on the degrees of exposure, and the PFAS concentrations in women’s follicular fluid have been shown to be consistent to that in the blood [20, 23, 25–30, 39, 43, 82]. During eIVFG, half of follicle culture media were replaced every other day and the conditioned follicle culture media were used for measuring the steroid hormone concentrations using ELISA. Follicles were imaged at each media change every other day using the 10x objective of the Olympus inverted microscope (Olympus Optical Co Ltd, Tokyo, Japan) for evaluating the follicle survival and size. Follicle death was defined by the morphological changes of oocytes with shrinkage, irregular shape, and/or fragmentation, and/or morphological changes of follicles with darker and/or disintegrated somatic cell layers. The follicle size was determined by averaging 2 perpendicular measurements from one side to another side of the theca externa per follicle using the ImageJ software (v1.53; National Institutes of Health, Bethesda, MD).

### Hormone assay

The concentrations of 17β-estradiol (E2), testosterone (T), and progesterone (P4) in the follicle culture media were measured using ELISA kits (Cayman Chemical, Ann Arbor, MI) according to manufacturer’s instructions. The mouse anti-rabbit IgG pre-coated wells were incubated with standards, conditioned culture media, rabbit antiserum, and hormone-acetylcholinesterase (AChE) conjugated for 60-120 min. After the wells were washed with washing buffer three times, the Ellman’s reagent was added and incubated at room temperature for 60-90 mins, and the absorbance was measured using a BioTek SpectraMax M3 microplate reader (BioTek Instruments, Inc., Winooski, VT) at 414 nm within 15 min. The inter-assay coefficients of variability were less than 15% and intra-assay coefficients of variability were less than 10% for all assays. The reportable ranges of E2, T, and P4 assay were 0.61-10,000, 3.9-500, and 7.8-1,000 pg/mL, respectively. N = 8 follicles for each group of treatments, each experiment was repeated for 3 times.

### *In situ* zymography

*In situ* zymography is a technique used to study activities of matrix metalloproteinases (MMPs) enzymes in fixed tissue samples. Follicles from day 6 of eIVFG were incubated in the growth media with 250 µM PFNA (N=8 follicles) or dimethyl sulfoxide (DMSO, vehicle control; N=8) for 2 hours, then transferred to the ovulation induction media containing 100 µg/mL fluorescent-conjugated DQ gelatin (Invitrogen) with the same concentration of PFNA and incubated at 37°C in a humidified atmosphere of 5% CO_2_ in air for 15 hours. After incubation, follicles were fixed with 3.8% paraformaldehyde (PFA) at 37°C for 1 hour, and then stained with DAPI and mounted for visualization. After proteolytic digestion by MMPs, the bright green fluorescence of the DQ gelatin is revealed and can be used to measure enzymatic activity. Fluorescent images of the DQ gelatin were obtained using EVOS cell imaging system (Thermo Fisher Scientific) at the excitation wavelength of 495 nm and detection wavelength of 515 nm.

### Single-follicle RNA sequencing (RNA-seq) analysis

RNA-seq was performed to investigate effects of PFNA on follicular transcriptomic profiling during FSH-stimulated follicle maturation or hCG-induced ovulation. Follicles were treated with vehicle (N = 10) or 250 µM PFNA (N=9) from either day 2 to 6 of eIVFG or for 4 hours during hCG-induced *in vitro* ovulation. Follicles at the end of either exposure window were collected, and total RNA were extracted using the Arcturus PicoPure RNA isolation kit (Applied Biosystems) according to manufacturer’s instructions. The library preparation and low-input RNA sequencing were performed on an Illumina NovaSeq PE150 platform by Novogene (Novogene Corporation, Sacramento, CA).

High-quality trimmed paired sequencing reads were uploaded into the Partek Flow software for RNA-seq data analyses. The potential rDNA and mtDNA contaminants were filtered using Bowtie 2. The filtered reads were aligned to the whole mouse genome assembly-mm10 using the HISAT 2 aligner. Raw read counts were obtained by quantifying aligned reads to Ensembl Transcripts release 99 using the Partek EM algorithm and then normalized based on the Transcripts Per Million (TPM) method. Pseudo genes were filtered out using the list of protein encoding genes from HUGO Gene Nomenclature Committee [83]. Principle component analysis (PCA) were performed using the PCAtools package [84]. Differential expression analysis was performed using the DEseq2(R) and genes with an absolute fold change >= 2.0 or <= 0.5 and a false discovery rate (FDR) adjusted *p*-value < 0.05 were defined as significantly differentially expressed genes (DEGs). Gene ontology (GO) analysis were performed using DAVID [85]. Kyoto Encyclopedia of Genes and Genomes (KEGG) pathway enrichment analysis were performed on cluster-associated genes using the WebGestalt [86].

### Mouse superovulation, PFNA exposure *in vivo*, and oocyte and ovary collection

An *in vivo* mouse superovulation model was used to investigate the effects of PFNA on gonadotropin-dependent follicle maturation and ovulation. Due to the fact that PFAS accumulate in ovaries and have long half-lives, the exposure window of PFNA covered both FSH-stimulated follicle maturation and LH/hCG-induce ovulation to recapitulate the real-world continuous exposure to PFAS in women. The administration route of PFAS via IP was selected based a previous study from Wang et al. which demonstrated that the short-term administration of PFAS through IP at 5 – 25 mg/kg was comparable to the bioaccumulation of PFAS observed in long-term human exposure through contaminated drinking water or occupational exposure [87]. The IP route was also considered less stressful and safer for rodents, particularly for repetitive exposure studies [88, 89]. In addition, the metabolic fates of administered compounds through IP are similar to oral administration, as compounds need to pass through the liver before distributing in other organs [89, 90]. Thus, the exposure route of IP and a dose range of 5 – 25 mg/kg were used in the *in vivo* PFAS exposure experiments.

Twenty-one-day-old CD-1 female mice were selected for the superovulation induction because: (1) the CD-1 mouse strain is one of the most common outbred strains used for studying female reproductive toxicology. It has been demonstrated that CD-1 mice are sensitive to ovarian toxic chemicals, such as doxorubicin (DOX), bisphenol-A (BPA), phthalate, methoxychlor (MXC), genistein, dioxin, and 2,3,7,8– Tetrachlorodibenzo-p-dioxin (TCDD), etc. [91–107]; (2) Pre-pubertal mice at 21-day-old are commonly used for the superovulation induction and oocyte retrieval for studying oocyte biology [108, 109], because their ovaries have a good number of early-antral follicles which are sensitive to exogenous gonadotropins for the induction of follicle maturation, ovulation, and resumption of oocyte meiosis I; (3) pre-pubertal mice have not yet developed a mature hypothalamus-pituitary-gonad (HPG) axis, which allows us to investigate the direct effects of PFAS on the ovary and exclude potential influences of PFAS on the hypothalamus and pituitary; and (4) the absence of ovarian cyclicity allows for a better control and synchronization of exogenous gonadotropin-stimulated follicle maturation and ovulation.

To investigate the effects of PFNA exposure on ovulation, mice were randomly assigned into different treatment groups and were treated with 1xPBS (N=15) or 5 (N=13), 15 (N=8), or 25 mg/kg (N=9) PFNA through IP injection daily for 5 days. On day 3, mice were IP injected with 5 IU of pregnant mare serum gonadotropin (PMSG, ProSpec, East Brunswick, NJ) to stimulate the maturation of early antral follicles to grow to the preovulatory stage to reach full maturation. Forty-six hours after PMSG injection, mice were IP injected with 5 IU of hCG (Sigma-Aldrich, St. Louis, MO) to induce ovulation. Mice were sacrificed at 14 hours post-hCG injection and oocytes were collected from the ampulla region of both sides of oviducts. The numbers of ovulated oocytes were counted and recorded, and ovaries were collected for histology.

To investigate the effects of PFNA on the expression of genes crucial for ovulation, 21-day-old CD-1 female mice received the same regimen of of PMSG and hCG treatments for ovulation induction as described above. Mice were randomly assigned into two groups and were treated with 1xPBS (N=5) or 25 mg/kg PFNA (N=5) through IP injection daily for 5 days. Mice were sacrificed at 4 hours post-hCG injection, and both sides of ovaries were collected. The follicular fluid of each ovary was collected in ultrapure water by poking the antral follicles under microscope. One ovary from each mouse was used for isolating large antral follicles and follicular somatic cells for RT-qPCR, and the other ovary was used for *in situ* RNA hybridization.

To determine the effects of pharmacological inhibition of PPARγ on PFNA-induced ovulation failure, 21-day-old CD-1 female mice received the same regimen of PMSG and hCG treatments as described above. Mice were randomly assigned into four groups and were treated with 1xPBS (N=8), 1 mg/kg PPARγ antagonist GW9662 (N=5), 25 mg/kg PFNA (N=8), or co-treated with 25 mg/kg PFNA and GW9662 (1 hour before PFNA, N=9), through IP injection. Mice were sacrificed at 14 hours post-hCG injection to count the numbers of ovulated oocytes in both sides of oviducts as described above.

### RNA extraction and RT-qPCR

For the *in vitro* exposure experiment, follicles collected on day 6 of eIVFG were used to examine the expression of genes related to follicle maturation and ovarian steroidogenesis, and follicles collected at 4 hours post-hCG were used to examine the expression of ovulatory genes using RT-qPCR. For *in vivo* exposure studies, somatic cells from four isolated large antral follicles in each mouse ovary (N=5 collected at 4 hours post-hCG) were pooled together. Total RNA of each follicle or pooled follicular somatic cells was extracted using the PicoPure RNA isolation kit (Thermo Fisher Scientific). Total RNA was then reversely transcribed into cDNA using the Superscript III First-Strand Synthesis System with random hexamer primers (Invitrogen) and stored at –80°C. RT-qPCR was performed in 384-well plate using the Power SYBR Green PCR Master Mix (Thermo Fisher Scientific) in a ViiA 7 Real-Time PCR System (Thermo Fisher Scientific).

RT-qPCR thermocycle was programmed for 10 min at 95 °C, followed by 40 cycles of 15 second (s) at 95 °C and 40 s at 60 °C, and ended with a melting curve stage to determine the specificity of primers. The relative gene expression levels of each gene were normalized by the expression of glyceraldehyde-3-phosphate dehydrogenase (*Gapdh*). The primer sequences of all examined genes were listed in table S1. N = 8 follicles per group of treatment. Each experiment was repeated for 3 times.

### Histology and *in situ* hybridization

Ovaries were fixed in 10% formalin overnight, embedded in paraffin, and serially sectioned at 5 µm. Ovarian sections were stained with hematoxylin and eosin (H&E, ThermoFisher Scientific) for histological examination of un-ruptured large antral follicles. *In situ* hybridization was performed using ovarian sections at 4 hours post-hCG to examine the expression of ovulatory genes using the RNAscope Multiplex Fluorescent Detection Kit V2 and the HybEZ Hybridization System (Advanced Cell Diagnostics, Inc., Newark, CA) following the manufacturer’s instructions. In brief, ovarian sections underwent pretreatment with heat, hydrogen peroxide, and protease before hybridization with target gene probes for tumor necrosis factor alpha-induced protein 6 (*Tnfaip6*), steroidogenic acute regulatory protein (*Star*), and prostaglandin-endoperoxide synthase 2 (*Ptgs2*). Subsequently, an HRP-based signal amplification system was applied to the target probes, followed by fluorescent dye labeling. The ovary sections were treated with DAPI (Maravai LifeSciences, San Diego, CA) and then imaged using a confocal microscope (Leica, Wetzlar, Germany).

### Analytical measurement of ovarian PFNA

Ovarian accumulation of PFNA was determined using mice treated with PBS (N=3) or 25 mg/kg PFNA (N=3) at 4 hours post-hCG injection. The vehicle, PFNA, and gonadotropin treatment regimen was the same as described above. The sample preparation and extraction protocol followed a previous study from Tatum-Gibbs et al with minor modifications [110]. In brief, one ovary from each vehicle or PFNA-treated mouse was fully homogenized in 100 μL ultrapure water, and the other ovary was placed in 100 μL ultrapure water to poke all PMSG-stimulated large antral follicles (∼ 20) to release the follicular fluid. For analytical sample preparation, the ovary homogenate or follicle fluid solution were added with 100 μL 0.1 M formic acid and 1 mL of acetonitrile (ACN) and then vigorously mixed for 15 min and centrifuged at 12,000 rpm for 15 min at the room temperature. Then, 500 μL of the supernatant was transferred to a HPLC sampling vial. Matrix-matched standard curves were prepared by externally spiking PFNA into the appropriate ovary matrix collected from untreated mice. The extraction method for standards was the same as aforementioned. The range of the standard curve was 0 – 100 ng/mg ovary weight. Before extraction, bile acid (BA) was added to all samples as an internal standard due to its similar retention time and molecular weight with PFNA. Then, 62.5 ng BA was spiked to each sample to monitor the extraction efficiency and sample loss during sample preparation and detection.

PFNA was measured using a Dionex UltiMate 3000 ultra-high-performance liquid chromatography (UHPLC) system (Thermo Fisher Scientific) coupled with a Q Exactive HF Hybrid Quadrupole-Orbitrap mass spectrometer equipped with an electrospray ionization (ESI) source (UHPLC-high resolution mass spectrometry or UHPLC-HRMS). Chromatographic separation was conducted on a LC C-18 column (50 × 3 mm, 1.7 μm, Phenomenex) at 30°C. The injection volume of samples was 5 μL. A mobile phase of H2O with 5 mM ammonium acetate (solvent A) and methanol with 5 mM ammonium acetate (solvent B) was used, following a 15-min linear gradient elution: 0/5, 5/85, 10/100, 13.5/100, 13.51/5, 15/5 (min/B%) at a flow rate of 350 μL/min. Nitrogen was used for all gas flows. Data was collected in negative ESI mode using parallel reaction monitoring (PRM) acquisition mode. Collision energy was set to 15 for the PFNA precursor ion of 462.9635. PFNA Product ions of 168.99, 218.99, and 418.97 were monitored for analyte identification and quantification. Data acquisition and processing was carried out with Thermo Xcalibur (v.4.0.27.19) software. A solvent blank (HPLC grade ACN) was carried out after 6 samples to monitor for any background contamination.

### Benchmark Dose Modeling (BMD)

The USEPA BMD Software (BMDS) tool (online version 2023.03.1) [111] was used to perform the frequentist BMD modeling and determine the point-of-departure (PoD) for endpoints examined in both *in vitro* and *in vivo* exposure models, including follicle growth, ovulation, oocyte meiosis I resumption, and hormonal secretion of E2, P4, and T for *in vitro* exposure to PFAS with significant toxicity, and ovulation for *in vivo* exposure to PFNA. For dichotomous data, such as failure of follicle rupture, ovulation, and oocyte meiosis I resumption, the 10% extra risk of these endpoints was set as the benchmark response (BMR_10_) level to derive the benchmark dose/concentration (BMD_10_ or BMC_10_) and the corresponding 95% lower confidence limit (BMDL_10_ or BMCL_10_). For continuous data, such as hormone secretion and follicle diameter, relative deviation of 10% from the background level was set as BMR, and both the constant and non-constant variance cases were explored. The default model selection and restriction were used: for dichotomous data, the *Dichotomous Hill*, *Gamma*, *Log-Logistic*, *Multistage* and *Weibull* models were run restricted, and the *Logistic*, *Log-Probit*, *Probit* and *Quantal Linear* models were run unrestricted; for continuous data, the *Exponential*, *Hill*, *Polynomial*, and *Power* models were run restricted, and the Linear model was run unrestricted. Selection of the best models followed the EPA recommended guideline to determine BMD_10_ or BMC_10_ and BMDL_10_ or BMCL_10_ [112]. *In vitro* to *in vivo* extrapolation was performed using an uncertainty factor of 100 based on possible interspecies differences between rodents and humans (10-fold) and interindividual differences (10-fold) in response to a toxicant [113, 114].

## Statistical analysis

Statistical analysis was performed using the GraphPad Prism version 9 (GraphPad Software, San Diego, CA). One-way ANOVA was performed for Normality test and homogeneity of variances then followed by Student’s t-test to compare numerical data of two treatment groups, including the expression of follicle maturation, ovulation related genes between vehicle and 250 μM PFNA treated follicles, and PFNA accumulation in the whole ovary and follicular fluid. One-way ANOVA followed by a Tukey’s multiple comparison test was used to compare numerical data of multiple treatment groups, including the results of follicle diameter, hormone concentration, *in vivo* ovulation and follicle counting, and expression of follicle maturation and ovulation related genes. Fisher’s exact test was performed to analyze the categorical data of *in vitro* follicle rupture and the first oocyte polar body extrusion. The data was reported as the mean value ± standard deviation (SD). For the hormone concentration, the data of measured concentrations were log transformed to address the skewed distribution of the hormonal data and to make the data distribution meet the condition of the parametric statistical analysis for one-way-ANOVA between different treatment groups. Statistical significance was defined as a *p*-value < 0.05.

## Results

### 1. A tiered *in vitro* toxicity testing of ovarian disrupting effects of long-chain PFAS

We first used our established 3D *in vitro* ovarian follicle culture system, eIVFG, to investigate the ovarian impacts of PFAS and toxic mechanisms involved. We selected three common long-chain PFAS (PFNA, PFOA, and PFOS) and three short-chain PFAS (PFBS, PFHpA, and GenX) that are increasingly used. A tiered toxicity testing pipeline was designed as shown in Figure. 1A. In Tier 1, immature mouse follicles were treated with various concentrations of PFAS during the FSH-stimulated follicle maturation window and the hCG-induced ovulation window. The concentrations of PFAS ranged from 1 to 250 μM to mimic the low, medium, and high exposure levels in humans. After *in vitro* ovulation induction, follicles were cultured for 2 days to allow for luteinization and progesterone secretion. Several key follicular events were examined, including follicle development, hormone secretion, ovulation, and oocyte meiosis. PFNA that has been understudied but may reach similar or even higher contamination levels (e.g., in the public drinking water systems and blood of residents in Paulsboro, New Jersey, USA) [115][116] and disrupted multiple key follicular events in Tier 1 was advanced to Tier 2 in which a specific window of exposure was used to distinguish which follicular functions were perturbed. In Tier 3, more advanced and in-depth molecular and mechanistic investigations were performed, and verification was conducted in an *in vivo* mouse model.

**Figure 1.**
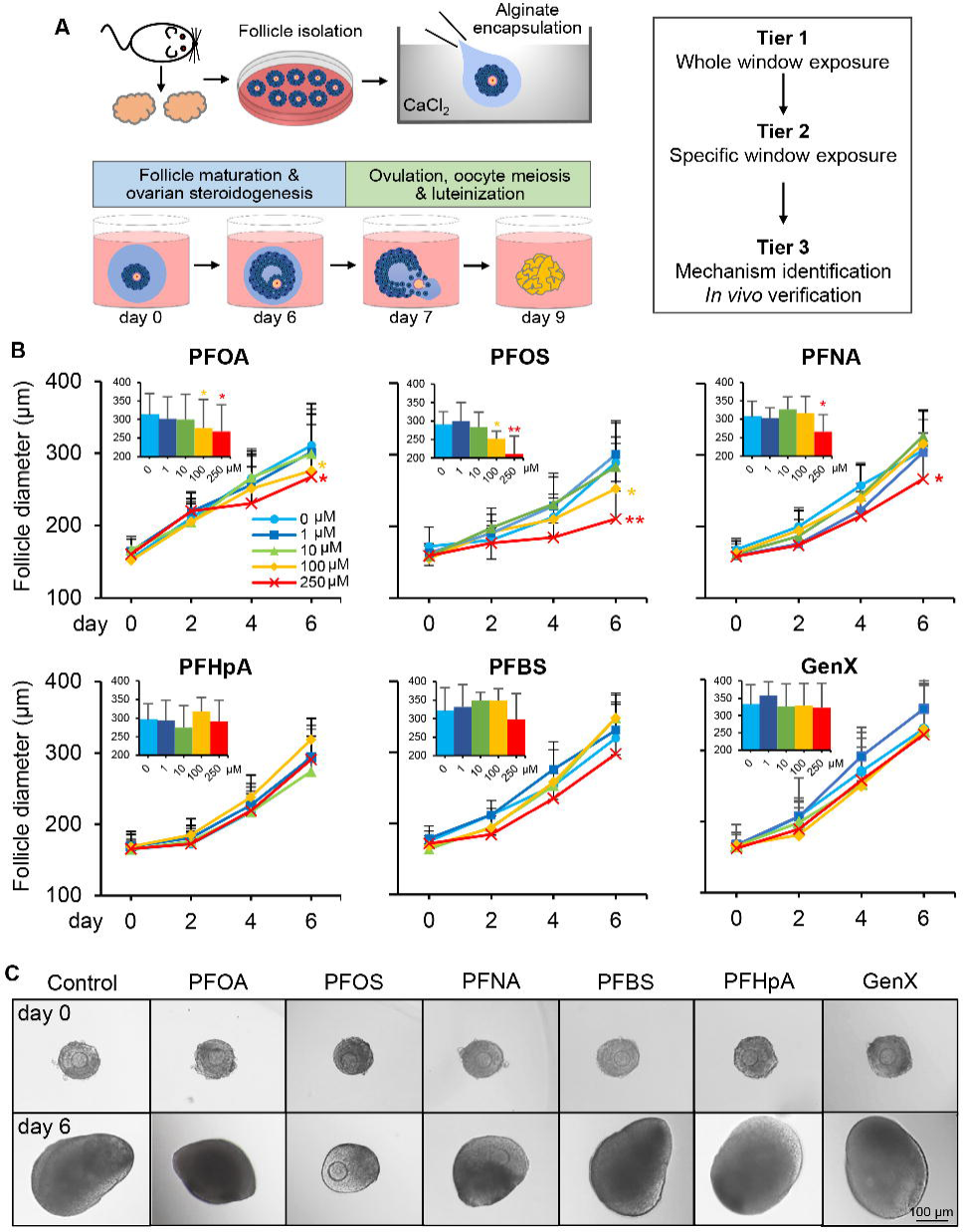
A tiered ovarian toxicity testing and the effects of PFAS on follicle growth. (A) The schematic of tiered ovarian toxicity testing starting from the 3D encapsulated *in vitro* follicle growth (eIVFG). (B) Effects of PFAS on follicle growth. Follicles were treated with either 0.1% DMSO as vehicle control, long-chain (PFOA, PFOS, and PFNA), or short-chain (PFHpA, PFBS, and GenX) at concentrations indicated from day 2-6 of eVIFG. Insets: Average follicle diameter on day 6. N=10-12 follicles in each treatment group. Data were analyzed using one-way ANOVA followed by a Tukey’s multiple comparisons test. (B). Shown are mean ± standard deviation; \**p*˂ 0.05 and \*\**p*˂0.01. (C) Representative follicle images on day 0 and 6 of eIVFG treated with either vehicle or PFAS as indicated.

Tier 1 testing showed that follicles treated with three long-chain PFAS at 1 and 10 μM and three short-chain PFAS at all concentrations were able to develop from the secondary stage to the antral stage with comparable size, morphology, and survival on day 6 of eIVFG; however, follicles exposed to 100 and 250 μM PFOA or PFOS or 250 μM PFNA had significantly smaller size (Figure. 1B and 1C). Follicles from day 6 were treated with hCG to induce ovulation. All three long-chain PFAS concentration-dependently inhibited follicle rupture, a key step of ovulation (Figure. 2A and 2B). Follicles treated with PFOA and PFNA at 250 μM and PFOS at 100 μM had significantly reduced percentages of ruptured follicles, and at 250 μM PFOS, no follicle ruptured (Figure 2A and 2B). The percentages of ovulated MII oocytes were concentration-dependently reduced by long-chain PFAS, with a decreasing trend and a borderline significance (*p*-value = 0.0769) for 250 μM PFOA, and no MII oocytes were ovulated from follicles treated with 250 μM PFOS (Figure. 2A and 2C). None of the three short-chain PFAS significantly affected follicle rupture or oocyte meiosis (Figure 2B and 2C, lower panels).

**Figure 2.**
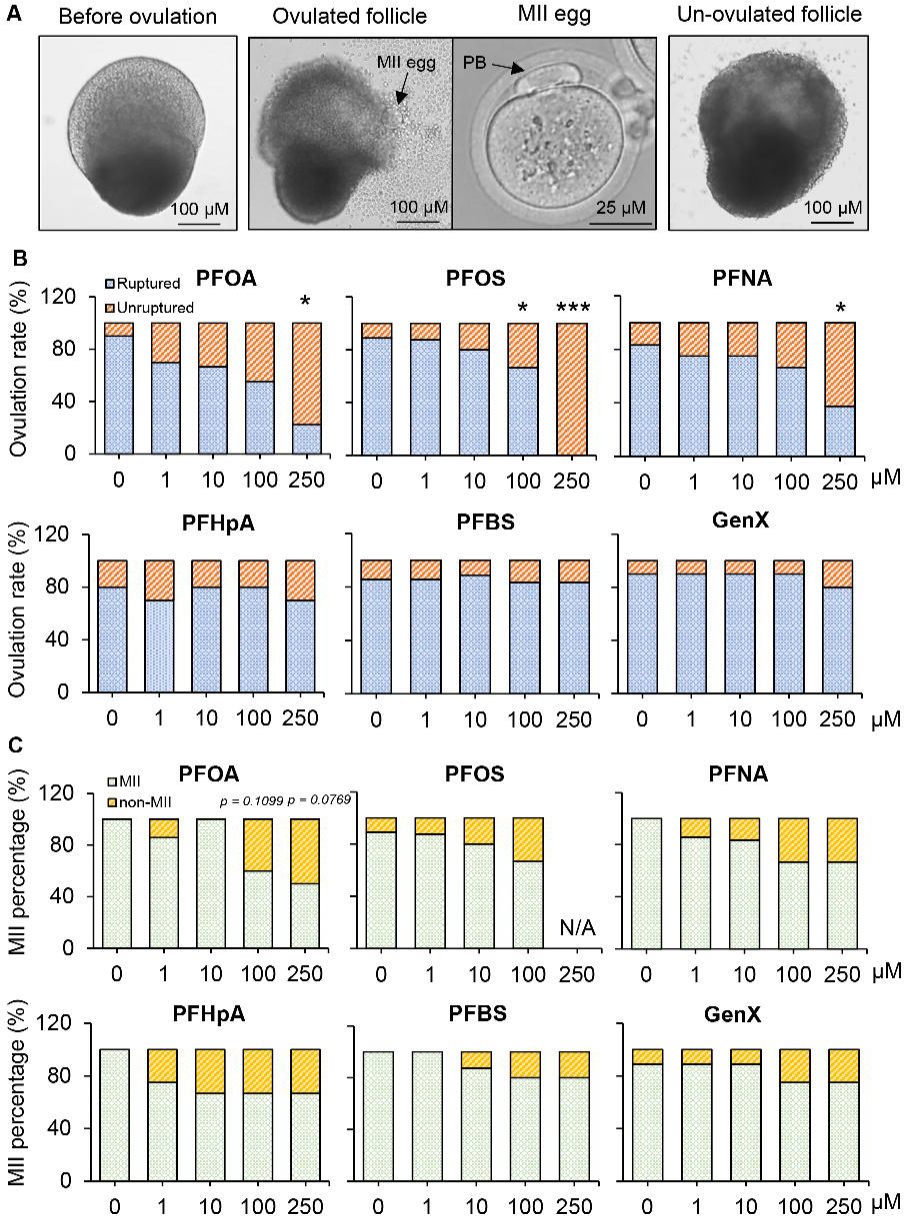
Effects of exposure to long-chain and short-chain PFAS during the entire gonadotropin-dependent follicle maturation and ovulation window on follicle ovulation. (A) Representative follicle images before and after hCG treatment, released oocytes at the meiosis of metaphase II (MII), and un-ruptured follicle after PFAS exposure. *In vitro* ovulation was induced by treating follicles with 1.5 IU/mL of hCG on day 6 of eIVFG for 14 hours. (B-C) Percentages of ruptured and un-ruptured follicles (B) and ovulated MII oocytes (C) treated with various concentrations of long– and short-chain PFAS. N=10 follicles in each treatment group. Statistical analysis was done by using Fisher’s exact test. \**p*˂ 0.05, \*\**p*˂0.01, and \*\*\**p*<0.001.

Ovarian steroidogenesis was next examined by measuring concentrations of E2 and T on day 6, corresponding to the end of the follicular phase of an ovarian cycle, and P4 on day 9 corresponding to the luteal phase. None of the three short-chain PFAS at any of tested concentrations affected the secretion of three hormones (Figure. 3). Follicles treated with 250 μM PFOA or PFOS had significantly lower concentrations of E2 (Fig. 3A) and T (Fig. 3B) on day 6, but they had comparable progesterone secretion to the control groups on day 9 (Figure. 3C). PFNA at 100 and 250 μM reduced the secretion of all three hormones. Although not all statistically significant, long-chain PFAS, particularly PFNA, at lower concentrations had the tendency to suppress ovarian steroidogenesis in a concentration-dependent manner. Altogether, these Tier 1 results indicate that long-but not short-chain PFAS exhibit ovarian disrupting effects by perturbing follicle growth, ovarian hormone secretion, and ovulation.

**Figure 3.**
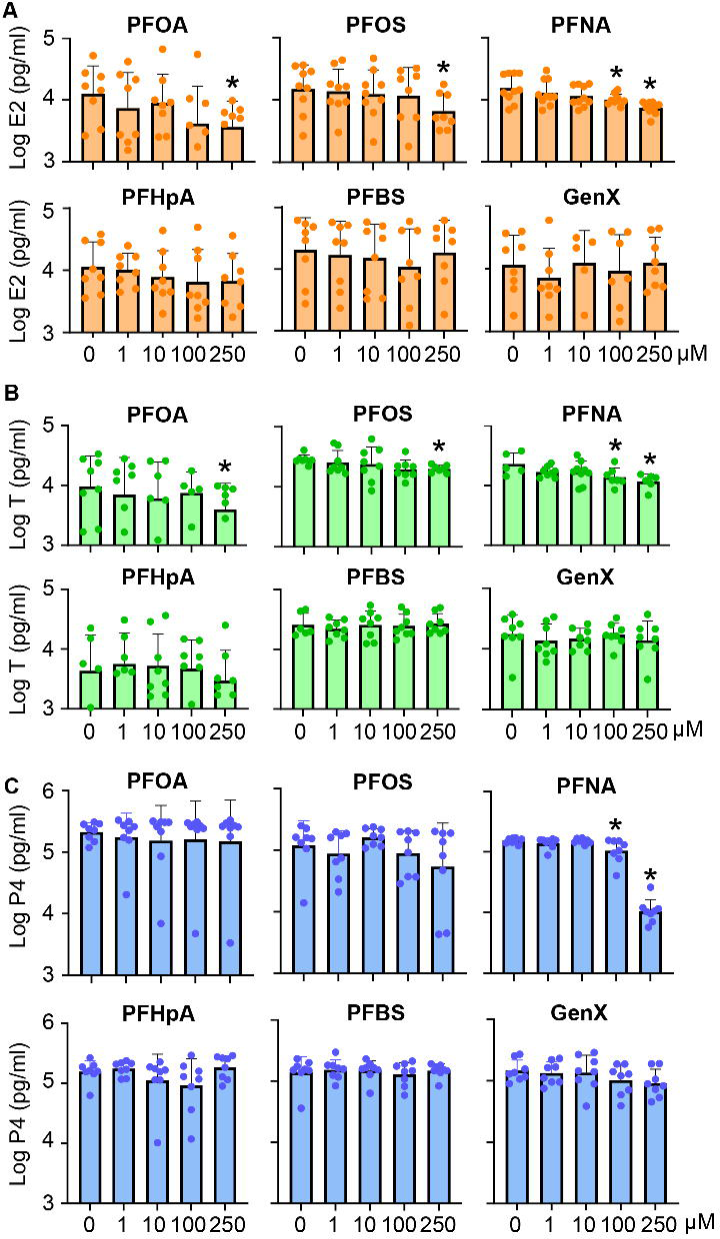
Effects of PFAS exposure during the entire gonadotropin-dependent follicle maturation and ovulation window on ovarian steroidogenesis. (A-B) Bars represent average Log_10_ concentration (pg/ml) of estradiol (A) and testosterone (B) in the conditioned follicle culture media collected on day 6 of eIVFG. (C) Average log_10_ concentration (pg/ml) of progesterone in the conditioned follicle culture media collected on day 9 after hCG-stimulated follicles were cultured for 48 hours. Data were analyzed with one-way ANOVA followed by a Tukey’s multiple comparisons test. N = 5-10 follicles in each group. Bars represent mean ± standard deviation; \**p*˂ 0.05 and \*\**p*˂0.01.

### 2. PFNA interferes with FSH-dependent follicle maturation to block ovulation

The ovulation failure and reduction of P4 from follicles treated with long-chain PFAS (Figure. 2 and 3) can result from three possibilities: (1) PFAS may affect FSH-induced follicle maturation toward the preovulatory stage, which causes ovulation failure as a secondary effect; (2) PFAS may directly compromise hCG-stimulated ovulation per se; or (3) both. In Tier 2, to distinguish the disrupting effects of PFAS on these two gonadotropin-dependent follicular events, we used PFNA to perform *in vitro* exposure that was restricted to either the follicle maturation window or the ovulation window. Short-chain GenX with the same exposure regimen was used as the negative control.

#### 2.1 PFNA alters the morphological and hormonal FSH-dependent follicle maturation

Follicles were first treated with the same concentration range of PFNA and GenX during the follicle maturation window from day 2 to 6 of eIVFG. Consistent to Tier 1, PFNA but not GenX concentration-dependently inhibited follicle growth and hormonal secretion of E2 and T (Figure. S1A). Upon hCG stimulation on day 6, PFNA at 1, 10, and 100 μM and all concentrations of GenX did not affect follicle rupture and oocyte meiosis on day 7 nor P4 secretion on day 9 (Figure. 4A-4C). However, 250 μM PFNA significantly reduced the percentage of ruptured follicles and P4 secretion except the percentages of ovulated MII oocytes (Figure. 4A-4C). Although we cannot rule out the possibility that the failed ovulation might be caused by the residual PFNA in follicular cells, as follicles were only treated with PFNA during the maturation window and there were suppressed follicle growth and ovarian steroidogenesis, these results strongly suggest that PFNA directly interferes with FSH-dependent follicle maturation to disturb subsequent ovulation and luteinization.

**Figure 4.**
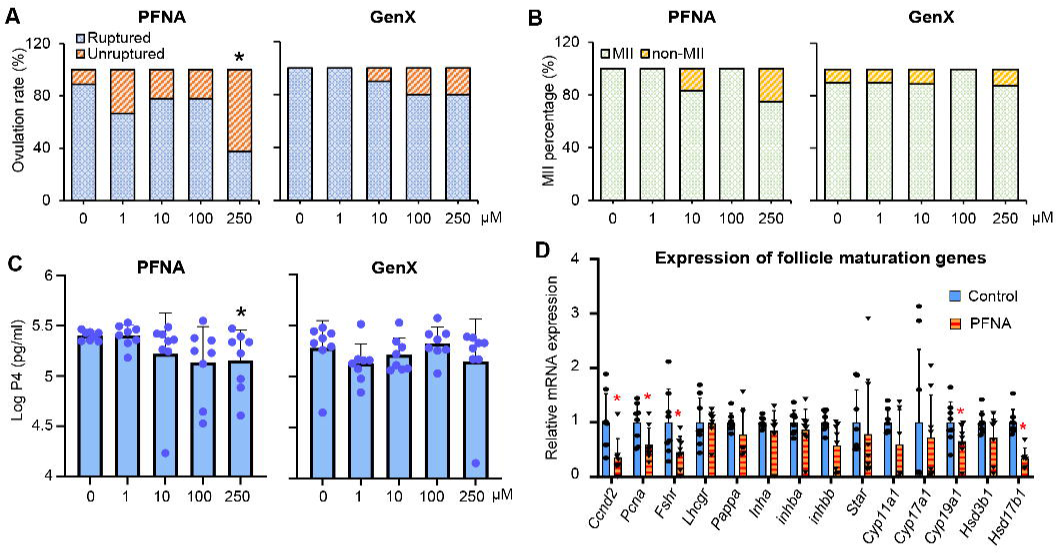
Effects of PFNA and GenX on follicle ovulation, resumption of oocyte meiosis, hormone secretion, and expression of follicle maturation genes. Follicles were exposed to various concentrations of PFNA or GenX from day 2 to 6 of eIVFG. (A-B) Percentages of ruptured and un-ruptured follicles (A) and ovulated MII oocytes (B) treated with various concentrations of PFNA or GenX. (C) Average Log_10_ concentration (pg/ml) of progesterone in the conditioned follicle culture media on day 9 after hCG-stimulated follicles were cultured for 48 hours. (D) Relative mRNA expression of follicle maturation genes examined by RT-qPCR in single follicles treated with 250 µM PFNA from day 2 to 6 of eIVFG. The mRNA expression levels were normalized by the expression of glyceraldehyde-3-phosphate dehydrogenase (*Gapdh*). Data were analyzed with student’s t-test. N = 8-10 follicles in each group. Shown are mean ± standard deviation; \**p*˂ 0.05 and \*\**p*˂0.01.

#### 2.2 PFNA alters the expression of FSH-induced follicle maturation genes

To identify the molecular targets of PFNA during the follicle maturation window, we performed a similar exposure experiment by treating follicles with vehicle or 250 μM PFNA from day 2 to 6 of eIVFG. Follicles were collected on day 6 for single-follicle qRT-PCR to examine the expression of several genes crucial for follicle maturation. The names, functions, and references of these genes were summarized in table S2. Results showed that PFNA significantly reduced the expression of cell proliferation genes, *Ccnd2* and *Pcna,* and other genes essential for granulosa cell differentiation and ovarian steroidogenesis, including *Fshr*, *Lhcgr*, *Cyp19a1*, and *Hsd17b1* (Figure. 4D). Although not statistically significant, PFAS had the tendency to decrease the expression of several other follicle maturation genes, including *Pappa*, *Inha*, *Inhba*, *Inhbb*, *Star*, *Cyp11a1*, and *Hsd3b1* (Figure. 4D). These results indicate that PFNA compromises follicle maturation by suppressing the expression of key FSH target genes.

#### 2.3 PFNA alters follicular transcriptome to perturb granulosa cell proliferation and differentiation

To better understand the effects of PFNA on follicular cell gene expression at the whole transcriptomic level, we collected follicles treated with vehicle or 250 μM PFNA using the same exposure regimen for single-follicle RNA-seq analysis. PCA showed that most PFNA-treated follicles were clearly separated from vehicle-treated follicles (Figure. 5A), suggesting marked alteration of the follicular transcriptome by PFNA exposure. There were 1004 DEGs with fold change >= 2 or <= 0.5 and FDR < 0.05, including 337 up– and 667 down-regulated genes in PFNA-treated follicles, with the top 10 genes in each direction highlighted in the volcano plot (Figure. 5B). The complete list of all DEGs was deposited to the Gene Expression Omnibus (GSE227267). Consistent with the qRT-PCR data in Figure. 4D, the expression of the same set of follicle maturation marker genes were also significantly reduced in the RNA-seq analysis (Figure. 5C), confirming that PFNA suppresses the expression of FSH target genes to disrupt follicle maturation and other follicle/oocyte reproductive outcomes.

**Figure 5.**
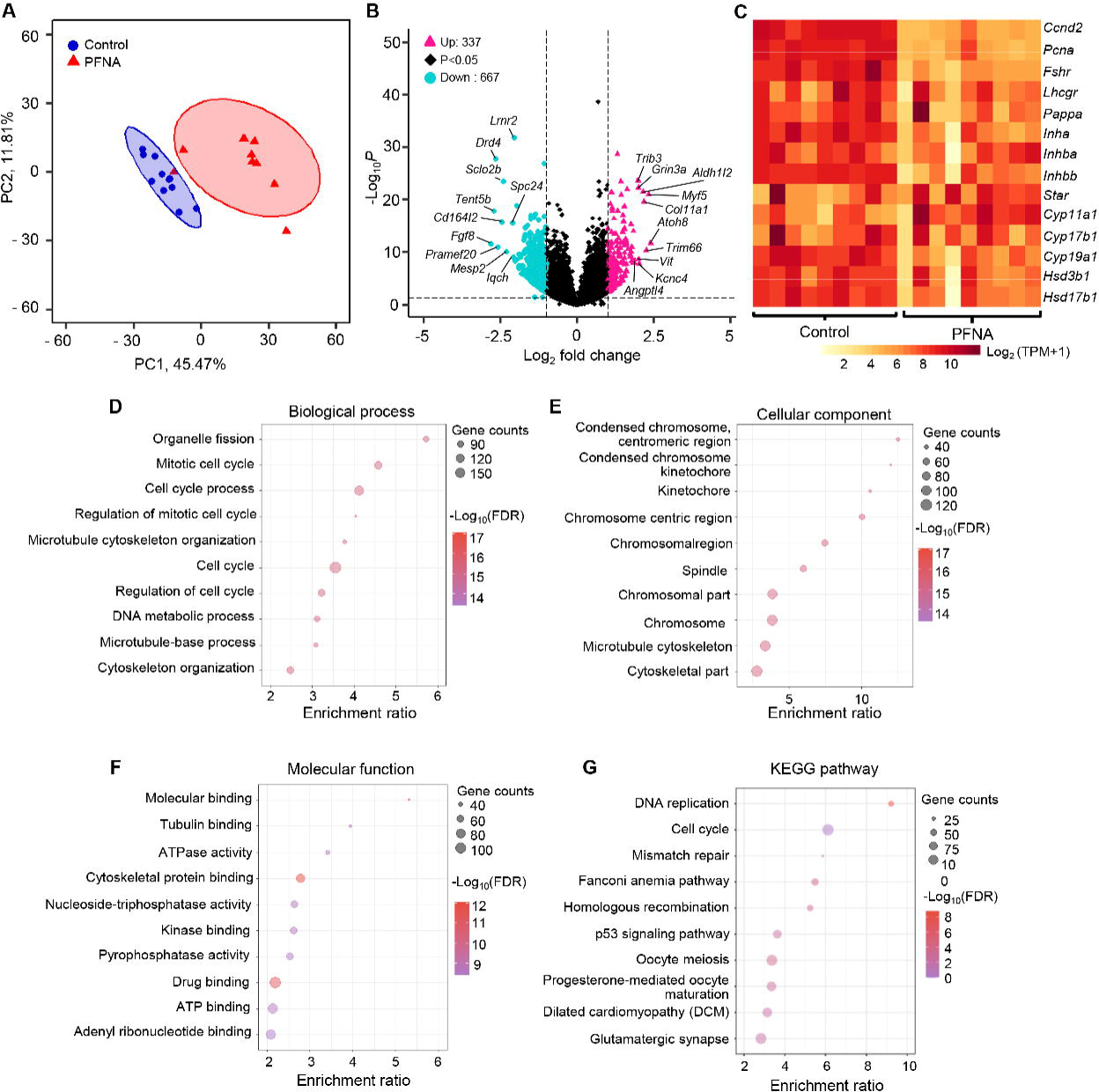
Single-follicle RNA-seq analysis reveals that PFNA exposure during the FSH-stimulated follicle maturation window alters follicular transcriptome. (A) Principal component analysis (PCA) of the first two PCs for follicles treated with PFNA at 250 μM (N=9) or vehicle (N=10). (B) Volcano plot of differentially expressed genes (DEGs, FDR < 0.05, absolute fold change >= 2 or <=0.5) in PFNA-treated follicles compared to the control. Pink-red: up-regulated genes; black: insignificantly altered genes; light-blue: down-regulated genes. (C) Heatmap of the same set of follicle maturation-related genes examined by both single-follicle RNA-seq (here) and qRT-PCR. Each column in the heatmap represents the relative change of expression level of genes for each sample. Gene ontology (GO) and KEGG pathway analysis of DEGs identified by single-follicle RNA-seq between follicles treated with PFNA at 250 µM (N=9) or vehicle (N=10) during the follicle maturation window. KEGG: Kyoto Encyclopedia of Genes and Genomes. (D-G) GO analyses of DEGs, including top 10 enriched biological processes (D), top 10 enriched cellular components (E), and top 10 enriched molecular functions (F). (G) Top 10 enriched KEGG pathways. Data represent in figure 5D-G are also presented in table S3.

DEGs were next used for the GO enrichment and KEGG pathway analyses. The specific up-/down-regulated genes for each enriched GO term and signaling pathway were listed in Table S3 and the top 10 GO terms and signaling pathways were highlighted in Figure. 5D-5G. Biological process analysis revealed that DEGs were primarily enriched in the processes of “Organelle fission” (ratio of up-/down-regulated genes, up/down:5/76), “Cell cycle” (up/down:14/159), and “Cytoskeleton organization” (up/down:14/75) (Figure. 5D). Cellular component analysis showed that DEGs were mainly related to “Condensed chromosome centromeric region” (up/down:1/37), “Kinetochore” (up/down:1/35), and “Spindle” (up/down:2/51) (Figure. 5E). Molecular function analysis showed that DEGs were closely associated with “Molecular binding” (up/down:14/65), “Tubulin binding” (up/down:6/30), and “ATPase activity” (up/down:5/35) (Figure. 5F). KEGG analysis revealed that several signaling pathways related to cell proliferation and DNA damage response (DDR) were significantly changed in PFNA-treated follicles, including “DNA replication” (up/down:0/15), “Cell cycle” (up/down:1/17), “Mismatch repair” (up/down:0/6), “Homologous recombination” (up/down:0/10), and “p53 signaling pathway” (up/down:2/10) (Figure. 5F). For instance, 5 out of 6 minichromosome maintenance protein complex (MCM) family genes (*Mcm 2-6*), the DNA ligase I gene (*Lig 1*), check point kinase 1 gene (*Chek1*), and DNA repair related genes (*Rad51*, *Rad51b*, *Rad54b*, and *Brca1*) were significantly reduced in PFNA-treated follicles. In addition, the pathways of “Oocyte meiosis” (up/down:1/17) and “Progesterone-mediated oocyte maturation” (up/down:1/13) were significantly enriched in the KEGG analysis (Figure. 5G). Together, these transcriptomic data are in line with the morphological results of PFNA-induced smaller follicle size and perturbed oocyte meiosis and ovarian steroidogenesis in both Tier 1 and 2 testing. The enrichment of “Cell cycle” and “DNA repair pathways” suggest that PFNA may perturb follicular cell proliferation and/or genome integrity.

#### 2.4 PFNA disrupts follicle maturation by activating PPARγ in granulosa cells

PFAS, particularly long-chain PFAS, have been shown to activate PPARs to produce toxicities [65, 66]. For the three PPAR subtypes, PPAR*α* and *β* are expressed in the ovarian stromal and theca cells, and their expression levels are stable through folliculogenesis and ovulation; whereas PPAR*γ* is primarily expressed in the mural granulosa cells of maturing follicles and is downregulated in preovulatory follicles and ovulating follicles [67–70]. Over-activating PPARγ with synthetic agonists such as troglitazone has been shown to inhibit granulosa cell proliferation [67, 70] and E2 secretion [67, 68, 70, 117–119] in a variety of species. These findings motivated us to hypothesize that PFNA acts as a PPARγ agonist to disrupt FSH-dependent follicle maturation and associated follicular functions.

To test this hypothesis, follicles were co-treated with 250 μM PFNA and various concentrations (0.1, 1, or 10 μM) of PPAR antagonists specifically targeting PPARα, β, or γ during the follicle maturation window. The three PPAR antagonists alone at all concentrations did not affect follicle growth and ovulation, and 250 μM PFNA alone consistently inhibited follicle growth and ovulation (Figure. 6A and 6B). PPARγ antagonist, GW9662, but not the PPARα (MK886) or PPARβ (GWK3787) antagonists, rescued the follicle growth and ovulation inhibited by PFNA (Figure. 6A and 6B). At hormonal levels, the antagonist targeting PPARγ but not PPARα or β, rescued PFNA-inhibited E2 and T secretion on day 6 (Figure. 6C). PPARα antagonist (MK886) appeared to rescue E2, but the result is not statistically significant (*p*-value=0.6649). At molecular levels, PPARγ antagonist significantly reversed the repression of follicle maturation-related genes in PFNA-treated follicles, including *Fshr*, *Cyp19a1*, *Hsd17b1*, and *Inhbb* (Figure. 6D). Collectively, these results demonstrate that PFNA acts as a PPARγ agonist as the MIE to interfere with FSH-dependent follicle maturation and associated follicular events.

**Figure 6.**
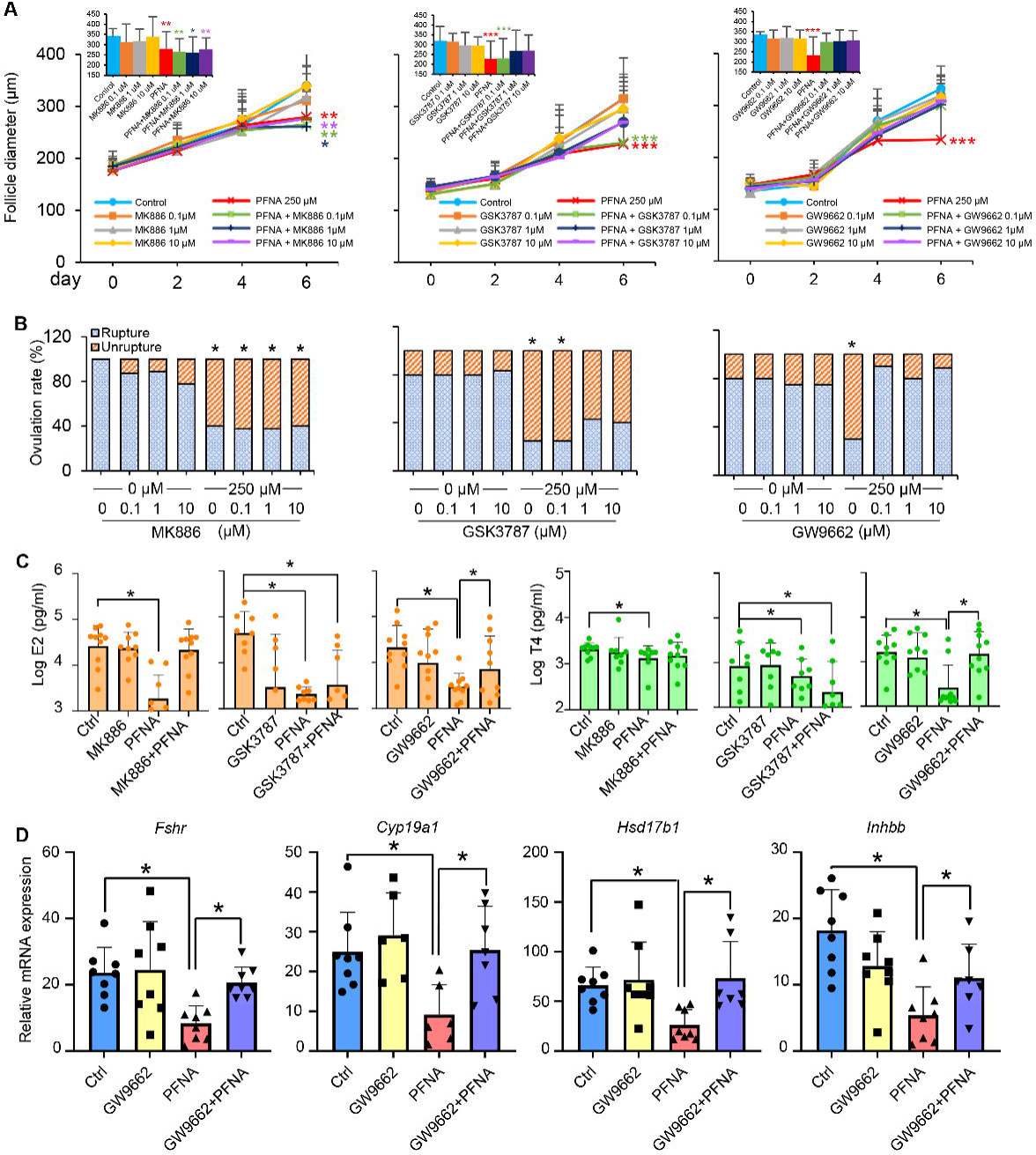
PFNA interferes with follicle maturation by acting as a PPARγ agonist. (A-B) Follicles were treated with 0, 0.1, 10 µM PPARα (MK886), PPARβ (GWK3787) or PPARγ (GW9662) antagonist as indicated, or 250 µM PFNA, or both 0.1-10 μM PPAR antagonists and 250 μM PFNA, or vehicle from day 4 to day 6 of eIVFG. (A) Average diameters of follicles. (B) Percentage of ruptured and un-ruptured follicles. (C-D) Follicles were treated with 10 µM PPAR antagonists or 250 µM PFNA alone or in combination of both, or vehicle control from day 4-6 of eIVFG. (C) Average log_10_ concentration of estradiol and testosterone in the conditioned follicle culture media collected on day 6. (D) Relative mRNA expression of follicle maturation genes examined by RT-qPCR (N=8) in follicles treated with vehicle, 10 μM PPARγ antagonist (GW9662), 250 μM PFNA, or both 10 μM PPARγ antagonist (GW9662) and 250 μM PFNA. Expression levels were normalized by the expression of *Gapdh*. Data were analyzed with student’s t-test (A), Fisher’s exact test (B), and one-way ANOVA followed by a Tukey’s multiple comparisons test (C and D), N=8-10 follicles in each treatment group. Shown are mean ± standard deviation; \**p*˂ 0.05 and \*\**p*˂0.01, and *\*\*\*p*<0.001.

### 3. PFNA inhibits LH/hCG-dependent follicle ovulation

To determine whether long-chain PFAS directly affect ovulation per se, similar to the PFAS exposure during the follicle maturation window, we chose PFNA and GenX to treat follicles with the same concentration range and only during the hCG-induced ovulation window. Follicles treated with PFNA at 1 and 10 μM and with GenX at all concentrations had comparable ovulation outcomes on day 7 and P4 secretion on day 9 (Figure. 7A-7C). However, 250 μM PFNA significantly inhibited follicle rupture and PFNA at 100 and 250 μM reduced P4 secretion (Figure. 7A and 7C). These results suggest that PFNA directly impacts ovulation and the subsequent luteinization and P4 secretion.

**Figure 7.**
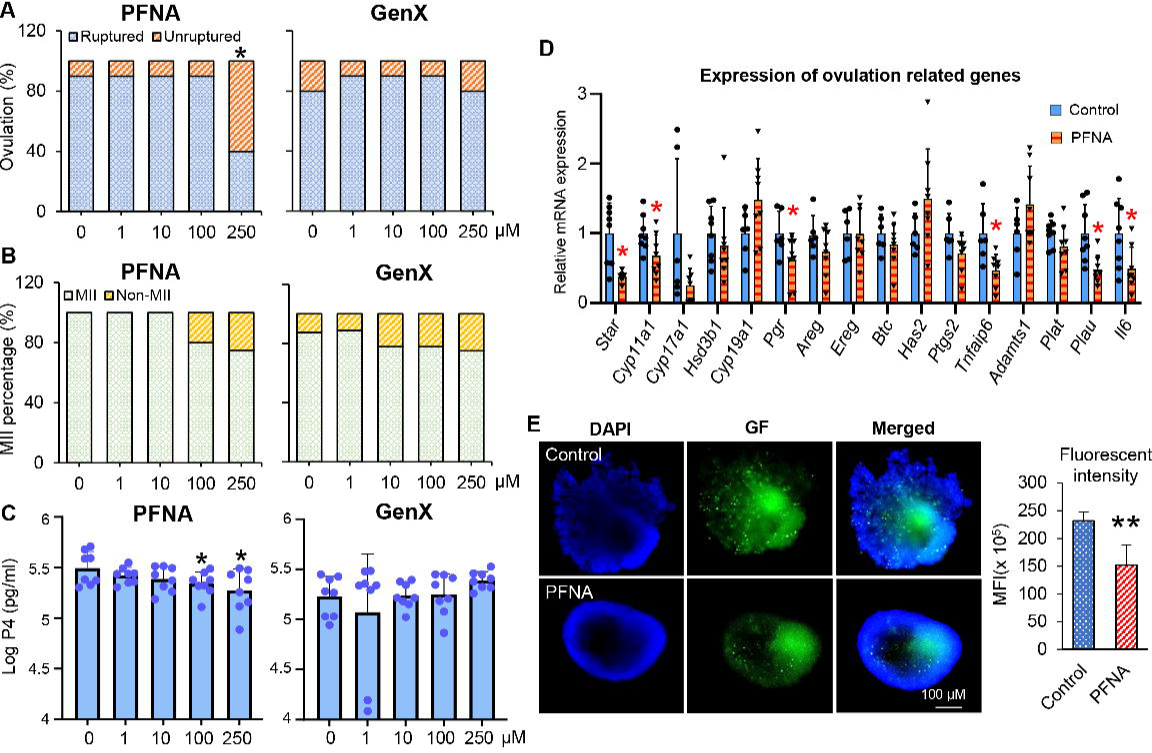
Effects of PFNA and GenX on follicle ovulation, expression of ovulatory genes, and activation of gelatinases. Follicles were exposed to various concentrations of vehicle, PFNA, or GenX as well as 1.5 IU/mL hCG on day 6 of eIVFG for *in vitro* ovulation induction. (A) Percentages of ruptured and un-ruptured follicles treated with various concentrations of PFNA and GenX. (B) Percentage of ovulated MII oocytes. (C) Average log_10_ concentration (pg/ml) of progesterone in the conditioned follicle culture media after hCG-stimulated follicles were cultured for 48 hours (N=8). (D) Relative mRNA expression of ovulation-related genes at 4 hours of post-hCG treatment examined by RT-qPCR. Expression data were normalized with expression of *Gapdh*. (E) Representative images and quantification of *in situ* zymography of follicles treated with vehicle or PFNA at 14 hours post-hCG; MFI: mean fluorescent intensity. Data were analyzed with Fisher’s exact test (A and B), and student’s t-test (C-E). Shown bars represent mean ± standard deviation; N=8-10 follicles in each treatment group. \**p*˂ 0.05 and **p˂0.01.

#### 3.1 PFNA alters the expression of key genes involved in follicle rupture and luteinization

Because the majority of LH/hCG target genes are highly induced at 4 hours in both *in vivo* and *in vitro* ovulation models [81, 120], we collected follicles at 4-hour post-hCG to examine the expression of several established ovulatory genes using single-follicle qRT-PCR. The names, functions, and related references are summarized in table S4. PFNA significantly reduced the expression of key ovulatory genes, including *Pgr*, *Tanfip6*, *Star*, *Cyp11a1*, *Plau*, and *Il6* (Figure. 7D). The reduction of *Star* and *Cyp11a1*, two genes involved in luteinization and P4 synthesis, is consistent with the decreased secretion of P4 in the formed CL organoids (Figure. 7C). PLAU has been shown to cleave plasminogen to active plasmin which further activates MMPs to degrade ECM components during ovulation [121, 122]. We collected follicles at 14-hour post-hCG to examine gelatinase (MMP2/9) activities using *in situ* zymography. Compared to the control group, follicles treated with PFNA had markedly reduced GFP fluorescent signals (Figure. 7E), indicating decreased gelatinase activation. This result is consistent with the reduced transcription of *Plau*, suggesting that the failed follicle rupture caused by PFNA might be due to diminished activation of gelatinase and associated ECM remodeling.

#### 3.2 Single-follicle RNA-seq analysis reveals perturbation of inflammatory and other signaling pathways during *in vitro* ovulation in PFNA-treated follicles

To identify molecules responsible for PFNA-induced ovulation failure in an unbiased manner, we collected follicles treated with vehicle or 250 μM PFNA for 4 hours for single-follicle RNA-seq analysis. PCA separated vehicle and PFNA-treated follicles into two distinct clusters (Figure. 8A), suggesting altered follicular transcriptome. There were 4193 significant DEGs with fold change >= 2 or <= 0.5 and FDR < 0.05, including 3103 up and 1090 down-regulated genes induced by PFNA during *in vitro* ovulation (Figure. 8B). The complete list of all was available at the Gene Expression Omnibus (GSE227267). The top 10 genes in each direction are highlighted in the volcano plot in Figure. 9B. Consistent with the qRT-PCR data (Figure. 8D), the expression of the same set of ovulatory genes was significantly reduced in the RNA-seq analysis (Figure. 8C).

**Figure 8.**
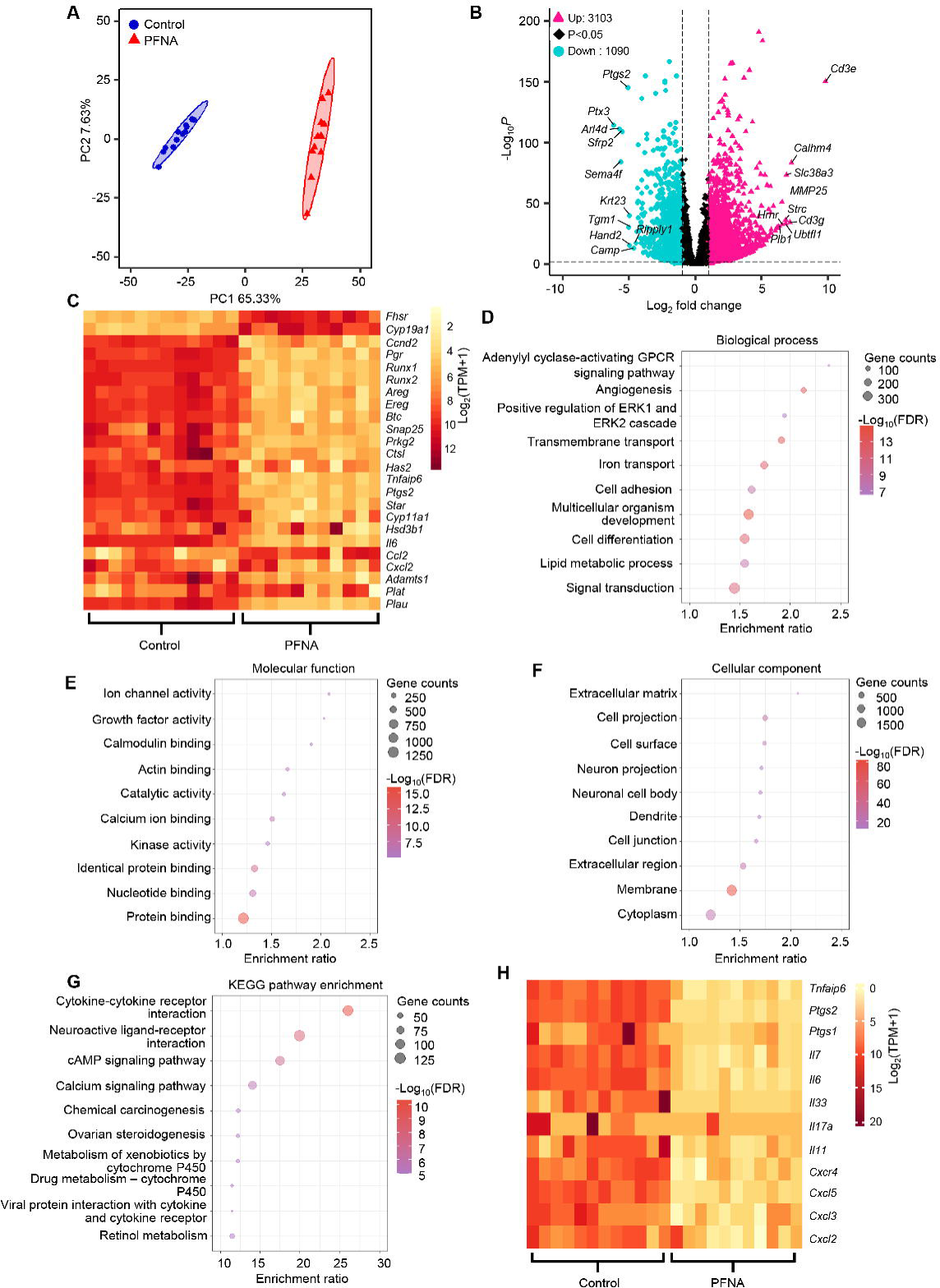
Single follicle RNA-seq analysis of follicles exposed to PFNA during ovulation window only. (A-H) Follicles were treated with vehicle control or 250 µM PFNA and 1.5 UI/ml hCG for 4 hours on day 8 of eIVFG. (A) PCA of the first two principal components for follicles treated with PFNA (N=10) or vehicle control (N=11). (B) Volcano plot of differentially expressed genes (DEGs, FDR < 0.05, absolute fold change > 2 or <-2) in PFNA-treated follicles compared to the control group. Pink red: up-regulated genes; black: non-significantly altered genes; light-blue: down-regulated genes. (C) Heatmap indicating relative change of ovulatory genes in PFNA-treated follicles (column 12-21) and control (column 1-11). (D-F) GO analyses of DEGs, including top 10 biological process enrichment results (D), top 10 cellular component enrichment results (E), and top 10 molecular function enrichment results (F). (G) Top 10 KEGG pathway enrichment results. (H) Heatmap indicating relative change of inflammatory genes in PFNA-treated follicles (column 12-21) and control (column 1-11). Data presented in figure D-G is also included in table S5.

**Figure 9.**
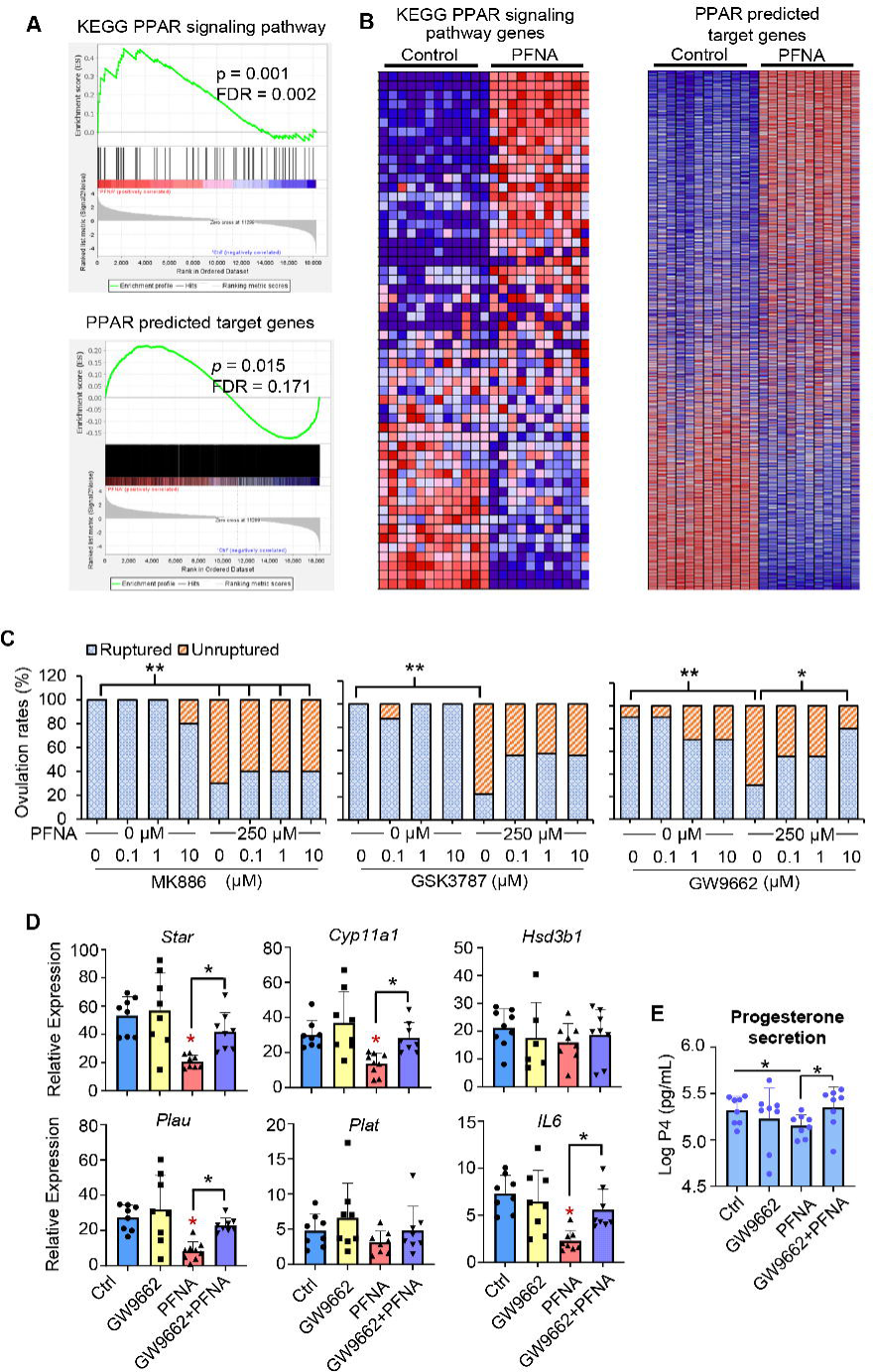
PFNA disrupts follicle maturation by activating PPARγ. (A) Gene set enrichment analysis (GSEA) of DEGs including KEGG PPAR signaling pathway gene sets and predicted PPAR target genes. (B) Heatmap of genes in the KEGG PPAR signaling pathway and predicted PPAR target genes. Columns represent the relative change of expression level of genes of each sample. (C) Follicles were treated with 0, 0.1, 1, 10 µM PPAR antagonists, or 250 µM PFNA, or combination of PPAR antagonists and PFNA, or vehicle control. Follicle rupture was then examined at 14-hours post-hCG and (N=10). (D) Relative mRNA expression of ovulatory genes examined by RT-qPCR; follicles were treated with 10 µM PPARγ antagonist (GW9662), or 250 µM PFNA, or combination of both, or vehicle control with hCG for 4 hours (N=8). The expression level of each gene was normalized by the expression of *Gapdh*. (E) Average log_10_ concentration of progesterone in conditioned culture media collected 48 hours post-hCG (N=8-10). Statistical analyses were done with fisher’s exact test (B) and one-way ANOVA followed by a Tukey’s multiple comparisons test (C and D). Bars represent mean ± standard deviation; \**p*˂ 0.05 and \*\**p*˂0.01, *\*\*\*p*<0.001.

DEGs were used for GO and KEGG pathway analyses. The specific up–/down– regulated genes for each enriched GO term and signaling pathway were listed in table S5. Biological process analysis showed that DEGs were primarily enriched in pathways related to “GPCR” (up/down:39/17), “Angiogenesis” (up/down:73/40), and “ERK1/2” (up/down:55/30) (Figure. 8D). Molecular function analysis revealed that DEGs were largely associated with “Ion channel” (up/down:60/8), “Growth factor activity” (up/down:33/26), “Calmodulin” (up/down:59/13), “Actin binding” (up/down:104/21), and “Catalytic activity” (up/down:86/36) (Figure. 8E). Cellular component analysis showed that DEGs were closely involved in “Extracellular matrix” (up/down:77/34), “Cell projection” (up/down:297/87), and “Cell surface” (up/down:159/92) (Figure. 8F). KEGG analysis identified the “Cytokine-cytokine receptor interaction” (up/down:63/37) as the most significantly altered signaling in PFNA-treated follicles, followed by the “cAMP signaling pathway” (up/down:66/26), “Calcium signaling pathway” (up/down:64/18), and “Ovarian steroidogenesis” (up/down:18/8) (Figure. 8G).

Ovulation has been proven as an inflammatory process [123]. LH stimulates the expression of proinflammatory factors at the early stage of ovulation, such as cytokines and prostaglandins to promote immune cell attraction, proteolysis, ECM remodeling, and angiogenesis [124, 125]. Our RNA-seq analysis showed that PFNA suppressed the expression of many inflammatory genes, including those encoding cytokines (*Il6*, *Il7*, *Il11*, *Il17a*, *Il33*, *Cxcl2*, *3*, 4, and *5*), cytokine receptors (*Cxcr4*), and other proinflammatory factors that crucially govern ovulation (*Ptgs1*, *Ptgs2*, and *Tnfaip6*) (Figure. 8H). These results demonstrate that PFNA interferes with the inflammatory response during hCG-induced ovulation to compromise follicle rupture.

We next compared PFNA-induced DEGs with another set of DEGs that we previously published at 4-hour post-hCG compared to 0-hour in follicles without any xenobiotic treatment in the same *in vitro* ovulation system [81]. There were 1521 LH/hCG target genes consistently changed between follicles treated with hCG and PFNA and follicles treated with hCG alone, suggesting adverse impacts of PFNA on the expression of LH/hCG-target genes (Figure. S2A and table S6.1). However, another 3921 non-LH/hCG target genes were selectively up– or down-regulated by PFNA (Figure. S2A and Table S6.2). KEGG pathway analysis revealed that those overlapped LH/hCG target genes were closely related to “Ovarian steroidogenesis”, “Peroxisome”, and “TNF signaling pathway” (Figure. S2B); and the non-overlapped genes were primarily associated with “Complement and coagulation cascades”, “Hematopoietic cell linage”, and “Aldosterone synthesis and secretion” (Figure. S2C). These results suggest other possible toxic effects and mechanisms of PFNA on follicle ovulation.

#### 3.3 PFNA activates PPARγ to inhibit ovulation and disrupt ovulatory signaling related to inflammation, proteolysis, and steroidogenesis

Activation of PPARγ has been shown to inhibit the expression of cytokines, such as IL-6 [71, 72]. A decrease of PPARγ activity is crucial for the induction of cytokines to underpin ovulation [126]. Since long-chain PFAS may activate PPAR to exhibit endocrine disrupting effects and our RNA-seq analysis revealed the suppression of many inflammatory genes in PFNA-treated follicles (Figure. 8H) [65], we hypothesize that PFNA activates PPARγ to interfere with inflammatory signaling and disrupt ovulation, particularly the rupture of ovulating follicles. To test this hypothesis, we first downloaded the gene list of PPAR pathway from KEGG and the predicted PPAR target genes from the PPAR gene database and performed Gene Set Enrichment analysis (GSEA). The PPAR pathway gene set in PFNA-treated follicles was significantly enriched when comparing to the control group, with the enrichment score of 0.45, *p*=0.001, and FDR = 0.002 (Figure. 9A). Predicted PPAR target gene set has the enrichment score of 0.22. This gene set is significantly enriched with *p*-value of 0.015 and FDR = 0.171 (Figure. 9A). The heatmap in Figure. 9B also showed that the expression of PPAR-associated genes and signaling in PFNA-treated follicles was significantly different from the control.

To further decipher the causation between PFNA-activated PPARγ and ovulation failure, follicles were co-treated with 250 μM PFNA and various concentrations (0, 0.1, 1, and 10μM) of PPAR antagonists targeting PPARα, β, and γ. Three PPAR antagonist alone at all concentrations did not affect ovulation (Figure. 9C). PFNA at 250 μM consistently inhibited follicle rupture (Figure. 9C), expression of ovulatory genes (Figure. 9D), and P4 secretion (Figure. 9E). The co-treatment of 10 μM PPARγ antagonist but not the antagonists of PPARα and PPARβ effectively and concentration-dependently rescued PFNA-induced follicle rupture failure (Figure. 9C). The co-treatment with the PPARβ antagonist (GSK3787) resulted in a partially rescuing effect in a non-concentration dependent manner, but the differences were insignificant in all three concentration groups with *p*-values of 0.0885, 0.1923, and 0.0885 for co-treatment of PFNA with 0.1, 1 and 10 µM GSK3787, respectively (Figure. 9C). At both molecular and hormonal levels, 10 μM PPARγ antagonist rescued the expression of key ovulatory genes inhibited by PFNA, including *Star*, *Cyp11a1*, *Plau*, and *Il6* (Figure. 9D) as well as P4 secretion in PFNA-treated follicles (Figure. 9E). Together, these results demonstrate that PFNA acts as an agonist of PPARγ in follicular granulosa cells to interfere with the induction of inflammatory factors and other ovulatory signaling, which eventually results in defective follicle rupture and P4 reduction.

### 4. PFNA inhibits gonadotropin-dependent follicle maturation and ovulation *in vivo*

We next used an *in vivo* mouse model to verify the ovarian disrupting effects and mechanisms of long-chain PFNA observed in the *in vitro* follicle culture system above as well as measure the ovarian accumulation of PFNA. As shown in Figure. 10A, CD-1 female mice at 21-day-old were treated with vehicle or 5, 15, and 25 mg/kg PFNA via IP for 5 days, with PMSG and hCG injections on day 3 and 5, respectively, to induce superovulation. This dosing regimen has been shown to result in a bioaccumulation of PFAS that is comparable to exposure to PFAS in humans through contaminated drinking water or occupational exposure [87]. The mouse ovulation results showed that PFNA-exposed mice had a significant and dose-dependent reduction of ovulated oocytes retrieved from both sides of oviducts compared with the control group, with 11.23 ± 8.22, 6.75 ± 7.27, and zero ovulated oocytes in mice treated with 5, 15, and 25 mg/kg PFNA, respectively, compared with 22.67 ± 11.59 oocytes in the control group (Figure. 10B). Ovarian histology and follicle counting results confirmed significantly more un-ruptured late-staged antral follicles in mice treated with 25 mg/kg PFNA (7.67 ± 2.16 in vehicle vs. 22.17 ± 2.40 PFNA-treated mice, Figure. 10C-10D).

**Figure 10.**
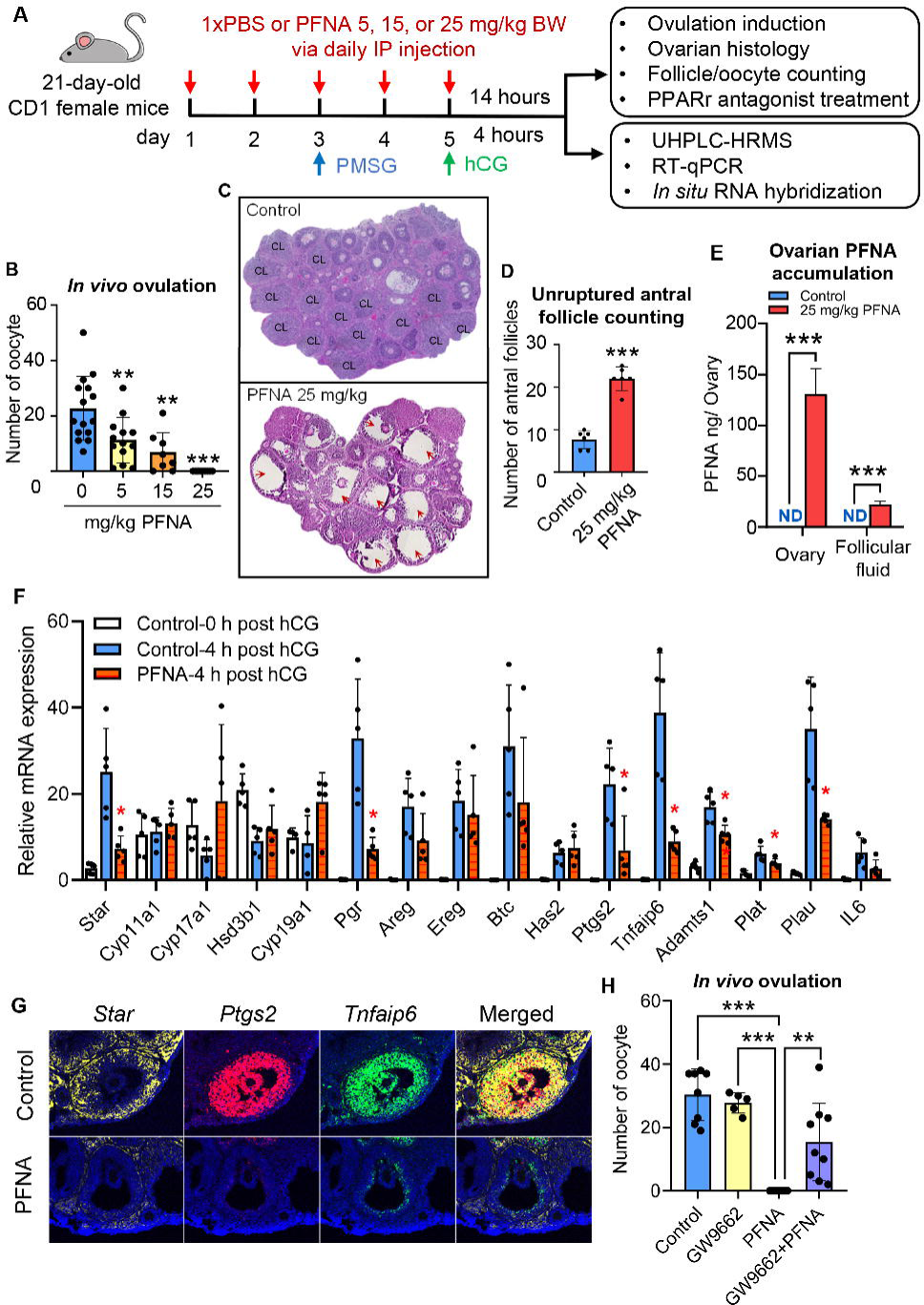
Effects of PFNA on ovulation in a mouse superovulation model. (A) Schematic of the superovulation model and PFNA exposure *in vivo*. (B) Average numbers of ovulated oocytes collected from mice treated with vehicle (N=15), 5 (N=13), 15 (N=8), and 25 (N=9) mg/kg of PFNA or PBS (control). (C) Representative images of mice treated with PBS and 25 mg/kg PFNA. Red arrows indicate un-ovulated late-stage antral follicles. (D) Average number of antral follicles in mice treated with PBS (N=5) or 25 mg/kg PFNA (N=5). (E) Analytical measurement of PFNA in the whole ovary and follicular fluid (N=3). ND: non detectable. (F) Relative mRNA expression of ovulation-related genes at 4 hours of post-hCG examined by RT-qPCR (N=5 in the control and N=5 in the PFNA treatment group). Expression data were normalized with *Gapdh*. (G) Representative images of *in situ* hybridization of large antral follicles treated with vehicle or PFNA at 4 hours post-hCG. (h) Average numbers of ovulated oocytes collected from mice treated with PBS (N=8), 1 mg/kg GW9662 (N=5), 25 mg/kg PFNA (N=8), or co-treatment of GW9662 and PFNA (N=9). Data were analyzed with student’s t-test (B and D). Error bars: mean ± standard deviation; \**p*˂ 0.05 and \*\**p*˂0.01.

The results of UHPLC-HRMS showed that PFNA were non-detectable in the ovary or the follicular fluid of vehicle-treated mice, indicating the minimum or negligible background contamination (Figure. 10E). PFNA accumulated in the ovary ( 130.7 ± 25.1 ng per ovary) and follicular fluid of large antral follicles ( 22.1 ± 3.1 ng per ovary) of 25 mg/kg PFNA-treated mice (Figure. 10E). The accumulated PFNA in the follicular fluid account for 17% of all PFNA in the ovary. Based on an estimation that ∼ 20 large antral follicles were pocked in each ovary to release the follicular fluid, and one antral follicle has the volume of 0.125 mm^3^ or μL (based on the diameter of 500 μm of an antral follicle and the weight of 6 mg per ovary), the PFNA concentration in the follicular fluid is 19.1 μM, which is comparable to PFAS concentrations in women’s systemic circulation and follicular fluid (1.5 to 220 µM) reported in previous studies [20, 23, 25–30].

We next collected follicular somatic cells from large antral follicles in the ovaries of PBS or 25 mg/kg PFNA-treated mice at 4 hours post-hCG injection for RT-qPCR and *in situ* RNA hybridization. Compared to the control group, PFNA significantly reduced the expression of ovulatory genes, including *Star*, *Pgr*, *Ptgs2*, *Tnfaip6*, *Adamts1*, *Plau*, and *Plat*, (Figure. 10F). *In situ* RNA hybridization results confirmed the transcriptional repression of *Star*, *Ptgs2*, and *Tnfaip6* in follicular theca and/or granulosa cells in the ovaries of PFNA-treated mice (Figure 10G).

To verify the mechanistic role of PPARγ in PFNA-induced defective follicle maturation and ovulation observed *in vitro*, 21-day-old CD-1 female mice were treated with vehicle, 25 mg/kg PFNA, 1 mg/kg PPARγ antagonist (GW9662), or both GW9662 and PFNA during PMSG/hCG-induced superovulation. Mice treated with vehicle or GW9662 alone had comparable numbers of ovulated oocytes (27.8 ± 3.28 in control vs. 34 ± 7.28 GW9662-treated mice, Figure. 10H). The treatment of 25 mg/kg PFNA alone consistently blocked ovulation; however, the co-treatment with GW9662 and PFNA resulted in a significant increase of ovulated oocytes (12 ± 10.12 in co-treatment vs. 0 in PFNA alone, Figure. 10H). Taken together, these *in vivo* exposure results demonstrate the bioaccumulation of PFNA in the ovary and follicular fluid; and at the phenotypical and molecular levels and mechanistic aspect, these *in vivo* results are consistent with PFNA-induced defective follicle maturation and ovulation observed *in vitro*.

### 5. BMD modeling for *in vitro* and *in vivo* endpoints

For the *in vitro* dual-window exposure (maturation + ovulation) experiments, the BMC_10_ and BMCL_10_ of endpoints that were significantly altered by PFAS are presented respectively in Supplemental Table S7, for continuous endpoints including follicle diameter, E2, T, and P4 secretion, and for dichotomous endpoints of follicle rupture and MII resumption. The BMC_10_ values range widely, between 2-2000 µM, while the BMCL_10_ values range between 1-160 µM. Extrapolating these values using an uncertainty factor of 100 for the *in vitro* to *in vivo* extrapolation suggests the reference concentrations of 10 nM – 1.60 µM for a 10% extra risk of follicular defects in humans. The BMCL_10_ for the long-chain PFOA, PFOS, and PFNA generally extend to a low concentration of 0.998 µM. The follicle rupture seems to be a more sensitive endpoint than hormone secretion and follicle growth, especially for the 3 long-chain PFAS. Interestingly, when the BMD analysis was applied to the distinct window exposure experiments for PFNA, the BMCL_10_ for inhibition of follicle rupture is lower when the exposure was limited to the maturation window (10.37 µM) than when it was limited to the ovulation window (28.83 µM) (Figure. 11A). Moreover, the BMCL_10_ of the dual-window exposure is 4.95 µM, suggesting that PFNA can more potently block follicle rupture when both maturation and ovulation processes are disrupted. In comparison, the oocyte meiosis I resumption does not seem to be sensitive to the exposure of either single or dual windows (Figure. 11B). For the *in vivo* superovulation experiment, the BMD_10_ and BMDL_10_ of PFNA for blocking ovulation were 0.91 and 0.56 mg/kg, respectively.

**Figure 11.**
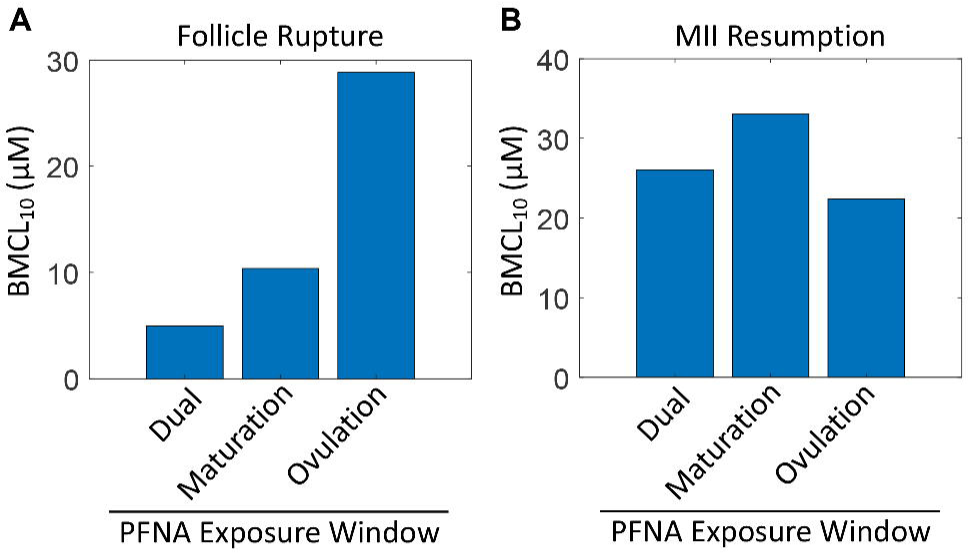
BMD modeling of in vitro and in vivo endpoints. (A) BMC_10_ and (B) BMCL_10_ of six PFAS species on dual-window exposure for *in vitro* endpoint as indicated. BMCL_10_ of PFNA for dual-, maturation-, and ovulation-window exposures as indicated for *in vitro* follicle rupture (C) and MII resumption (D). Data presented in this figure is also included in table S7.

## Discussion

Approximately 10-15% of reproductive aged women experience reproductive disorders and infertility, with ovarian disorders as the leading cause [127–129]. The etiology of these ovarian dysfunctions remain elusive but has been related to exposure to environmental EDCs [130], including PFAS [131]. PFOA and PFOS have been classified as Persistent Organic Pollutants (POPs) by the Stockholm Convention, an international environmental treaty, in 2017 [132], and both were designated as the hazardous substances by the US EPA [133]. Many countries globally have implemented laws and policies to reduce PFAS pollution and exposure [120]. Recently, the US EPA has joined these efforts by proposing to establish legally enforceable Maximum Contaminant Levels (MCLs) of six PFAS (PFOA, PFOS, PFNA, PFHxS, PFBS, and GenX) in drinking water through the National Primary Drinking Water Regulation (NPDWR) [134]. In line with these efforts, a decline in the blood levels of PFAS, particularly PFOA/S, has been observed in populations in the US [135, 136] and other countries [26, 137]. However, due to the high persistence and long half-lives, the long-chain legacy PFAS remains prevalent; moreover, the female reproductive impact of other long-chain PFAS such as PFNA and emerging short-chain alternatives, such as GenX, is largely unknown. Here, using a 3D *in vitro* ovarian follicle culture system and an *in vivo* mouse model, we demonstrated that long-chain PFAS at environmentally relevant exposure levels interfere with gonadotropin-dependent follicle maturation, ovulation, and hormone secretion; PFNA, an understudied long-chain PFAS but has been reported to reach similar or even higher contamination levels than the legacy PFOA/S in some community water bodies [138, 139], activate PPARγ in granulosa cells as the MIE to result in aforementioned ovarian disrupting effects.

Both of our *in vitro* and *in vivo* results demonstrate that long-chain PFAS interfere with ovulation. However, as the exposure of PFAS spanned both windows of follicle maturation and ovulation, the ovulation failure might result from either defective follicle maturation, direct impact of PFAS on ovulation, or both. Through two distinct *in vitro* PFNA exposures and in-depth analysis using eIVFG, long-chain PFAS, particularly PFNA, were found to perturb both gonadotropin-dependent follicular events. During the FSH-stimulated follicle maturation, PFNA delayed follicle growth and reduced hormonal secretion of E2 and T. In line with these morphological and hormonal changes, molecular analysis using RT-qPCR and RNA-seq revealed that PFNA suppressed the expression of key genes regulating granulosa cell proliferation (*Ccdn2* and *Pcna*), ovarian steroidogenesis (*Cyp19a1*, *Hsd3b1*, and *Hsd17b1*), and follicle maturation (*Fshr*, *Pappa*, and *Lhcgr*). Collectively, these results demonstrate that long-chain PFAS can interfere with ovarian follicle maturation, and the subsequent failure in ovulation, luteinization, and progesterone secretion could be a secondary consequence of the defective maturation.

When a mature follicle was exposed to PFNA only during the ovulation window, PFNA inhibited follicle rupture and P4 secretion. Moreover, PFNA suppressed the expression of key ovulatory genes involved in ECM remodeling (*Pgr*, *Plau*, and *Adamts1*), cumulus cell expansion (*Has2* and *Tnfaip6*), inflammation (*Il6*, *Cxcl2*, and *Ptgs2*), and luteal steroidogenesis (*Star*, *Cyp11a1*, and *Hsd3b1*). These findings demonstrate that long-chain PFAS also directly affect ovulation per se. As demonstrated by the BMD modeling analysis, when both the maturation and ovulation windows are perturbed by PFNA, the BMCL_10_ becomes much lower, indicating PFNA acts on the two processes synergistically to disrupt ovulation. Taken together, these experiments and analysis demonstrate that long-chain PFAS, particularly PFNA, can disrupt both gonadotropin-dependent follicle maturation and ovulation and associated ovarian steroidogenesis.

Accumulating evidence reveals that PFAS, particularly long-chain PFAS, can act as a PPAR agonist to exhibit toxicities, including endocrine-disrupting effects [65, 66]. PPARs are a family of nuclear transcription factors involved in various physiological and pathophysiological processes, such as lipid metabolism, inflammation, tissue remodeling, and cell proliferation [140, 141]. PPARs heterodimerize with 9-cis-retinoic acid receptors (RXRs) to form the PPAR/RXR heterodimer, which binds to the PPAR response element (PPRE) in the promoter region of PPAR target genes to regulate their transcription [142]. Although the exact functions of PPARs in the ovary are not fully understood, PPARγ has been shown to regulate granulosa cell proliferation, ovarian steroidogenesis, and tissue remodeling [143]. For instance, the overactivation of PPARγ through synthetic agonists (e.g., troglitazone) has been found to inhibit granulosa cell proliferation [67, 70], aromatase expression [117–119], and E2 secretion [67, 68, 70, 117–119, 144, 145]. In addition, the active metabolites of phthalates, such as mono-(2-ethylhexyl) phthalate (MEHP), have been found to activate PPARγ to inhibit aromatase expression and E2 secretion in both mouse primary granulosa cells [144, 146] and immortalized human KGN granulosa cells [147]. In response to the LH surge, the downregulation of the abundance and activity of PPARγ is obligatory for the induction of cytokines and other inflammatory factors to induced ovulation [126]. Our studies revealed that the selective antagonist targeting PPAR*γ* but not α and β effectively reversed PFNA-induced smaller follicle size, inhibited ovarian steroidogenesis, disrupted ovulation, and suppression of FSH/LH target genes. Moreover, the rescuing effects of PPAR*γ* antagonist on PFNA-induced follicle ovulation failure can be validated *in vivo*. These results suggest that PFNA acts as a selective PPARγ agonist in granulosa cells to interfere with gonadotropin-dependent follicle maturation and ovulation. During FSH-induced follicle maturation, PPARγ has been suggested as a transcriptional repressor to suppress the overexpression of LH-target genes, which prevents premature luteinization of granulosa cells [71, 72]. Our RNA-seq analysis revealed that follicles treated with PFNA during the follicle maturation window had significantly higher expression of several established LH-target genes, such as *Areg*, *Ereg*, *Pgr*, *Runx1/2*, *Il6*, and *Cxcl1* (Figure S3), which further confirms our hypothesis that PPARγ is the molecular target of PFNA and that exposure to PFNA may disrupt granulosa cell differentiation in maturing follicles.

The phase-out of legacy long-chain PFAS make short-chain PFAS increasingly manufactured and applied as alternatives [2, 3]. There are currently no regulatory guidelines regarding the use and safety levels of short-chain PFAS, and their impacts on female reproductive health have not yet been studied. Compared to long-chain PFAS, short-chain PFAS may have faster elimination rates and shorter half-lives [22, 64]. A recent study reported that short-chain PFAS had weaker binding affinity toward PPARs, which may result in less bioaccumulation and toxicities [65]. Another *in vivo* study conducted by the US National Toxicology Program (NTP) showed that in young adult female rats orally exposed to long-chain PFOS (C8) or short-chain PFBS (C4) for 28 days, PFBS required a much higher dose (250-1000 mg/kg) to disrupt rat estrous cyclicities to an extent similar to PFOS (5 mg/kg) [149]. Our results are in line with these findings and revealed that all three short-chain PFAS did not affect follicle reproductive outcomes as the long-chain PFAS did. However, we are cautious to conclude that short-chain PFAS are safer for female reproduction giving the following reasons. First, compared to long-chain PFAS, short-chain PFAS have a lower adsorption potential, making them highly mobile [150, 151], more prone to reach and contaminate water sources, and more difficult to be removed by large scale filtration system [152–154]. Second, short-chain PFAS might be as persistent as long-chain PFAS [155], leading to an increase in their environmental distribution and accumulation due to the growing use of short-chain PFAS as substitutes. Third, while short-chain PFAS have shorter half-lives after absorption, their environmental persistence and high contamination levels may result in continuous and chronic exposures in humans, raising concerns about the potentially harmful health impacts of the long-term exposure. Fourth, short-chain PFAS have been found to have a high affinity for binding to serum albumin [156, 157], which could lead to a high distribution and concentration in highly vascularized tissues, including antral follicles. Last but not least, short-chain PFAS have been shown to exhibit similar or even more toxic effects than long-chain PFAS in non-reproductive tissues through similar or distinct toxic mechanisms [75, 158, 159]. These facts highlight the need of further in-depth studies to fully understand their female reproductive impact.

Despite our *in vitro* exposure data showed that all three long-chain but not short-chain PFAS exhibited ovarian toxicities, but they had distinct toxic potencies and patterns on different follicular endpoints. The chemical structure of PFAS, especially the length of carbon backbone chains and functional groups attached, has been shown to determine the toxicity of different PFAS congeners [160]. Herein, we advanced PFNA for mechanistic assessments and *in vivo* verification but did not perform similar studies for PFOA/S and short-chain PFAS. Moreover, previous studies reported discrepancies of PPARs between mouse and humans, such as the less sensitivities in humans to PPARα activators due to a lower PPARα expression and DNA binding affinity compared to rodents, and different metabolic features such as the glucose-insulin singling pathway between mouse and human PPARγ mutants [161–163]. For instance, PPARα activators have been shown to induce liver cancer in mice by stimulating hepatocyte proliferation through a PPARα-dependent induction of MYC Proto-Oncogene, BHLH Transcription Factor (MYC) and cytotoxicity; in contrast, these effects were either absent or significantly ameliorated in similarly treated PPARα-humanized mice [164, 165]. With respect to the ovarian disrupting effects of short-chain PFAS, it is worth noting that while there were no significant morphological or hormonal changes in follicles treated with all concentrations, we cannot rule out the possibility that short-chain PFAS, such as GenX, did not perturb follicular health at molecular levels. Collectively, advanced investigations regarding the roles of different PPAR isoforms, structurally different PFAS congeners, possible perturbation of short-chain PFAS at molecular levels, and species difference in PFAS-induced ovarian toxicity are needed in future studies.

In line with folliculogenesis, the follicle-enclosed oocyte produces and accumulates maternal factors, such as mRNAs, proteins, and organelles, to acquire both meiotic and developmental competence to enable subsequent fertilization and early embryogenesis. Successful oogenesis relies on the bidirectional communications between oocytes and surrounding somatic cells [166, 167]. *In vitro* exposure of denuded oocytes to PFNA has been shown to affect oocyte maturation by inducing oxidative stress [168], indicating that PFAS may directly affect oocytes. Our results showed that there was a significant reduction of the percentages of ovulated MII oocytes released from PFAS-treated follicles. However, it is unclear whether this oocyte meiotic disorder is a direct perturbation of PFAS on oocytes, an indirect effect through granulosa cells, or both. In addition to somatic granulosa cells, PPARγ has also been detected in mammalian oocytes [117, 169]. Thus, future studies should determine whether PFAS can accumulate in oocytes and compromise oocyte quality by interfering with oogenic PPARγ and related signaling.

The results obtained from both an *in vitro* ovarian follicle culture system and an *in vivo* mouse model here suggested the ovarian disrupting effects of long-chain PFAS and the mechanisms via PPARγ. However, it is important to acknowledge several limitations needed to be addressed in future studies. First, the follicle culture system used a constant FSH concentration, which does not recapitulate the dynamic secretion of gonadotropins and lacks the negative and positive feedback control of the hypothalamus-pituitary-gonad (HPG) axis. Second, the regulatory signals and gene expression patterns involved in follicle development and ovulation are complex. Thus, the ratio of up-/down-regulated DEGs may not entirely reflect the activation/deactivation status of the corresponding enriched process or pathway in the PFNA-treated follicles, warranting further investigation in the future. Third, we primarily focused on the ovarian impacts of a single PFAS, but humans are exposed to PFAS in mixtures. Fourth, the mouse superovulation model and PFAS exposure via IP do not reflect the real-world exposure scenario of PFAS in humans. Thus, a long-term oral exposure (e.g., via drinking water) to PFAS and mixtures in a naturally cycling mouse model need to be considered in future studies.

In conclusion, we demonstrate ovarian disrupting effects of PFAS. PFNA, a long-chain PFAS that can have similar environmental contamination levels to the legacy PFOA/S, accumulates in the ovary and acts as a PPARγ agonist in follicular granulosa cells to interfere with follicle maturation, ovulation, and hormone secretion. The widespread contamination, bioaccumulation, and long half-lives of PFAS pose a potential risk to women’s reproductive health, including imbalanced ovarian hormone secretion, anovulation, irregular menstrual cycles, and infertility. There is an urgent need to reduce or eliminate exposure to PFAS to safeguard women’s reproductive health and fertility.

## Authors’ roles

P. Pattarawat and T. Zhan contributed to the experimental design, data collection and analysis, and manuscript writing. Y. Fan contributed to the statistics analysis and manuscript writing. J. Zhang, E. Kim, and Y. Zhang contributed to *in vitro* exposure data collection and hormone measurement. T. Zhan, H. Yang, and B. Buckley contributed to the analytical measurement of PFNA using HPLC-HRMS. NC. Douglas, M. Urbanek, J. Burdette, and JJ. Kim contributed to data interpretation and manuscript writing. S. Moyd contributed to BMD modeling and manuscript editing. Q. Zhang contributed to the experimental design, BMD modeling, and manuscript writing. S. Xiao conceived of the project, designed experiments, analyzed and interpreted data, wrote the manuscript, and provided final approval of the manuscript.

## Funding and acknowledgments

This work was supported by the National Institutes of Health (NIH) K01ES030014 and P30ES005022 to S. Xiao; R01ES032144 to S. Xiao and Q. Zhang; UH3ES029073 to M. Urbanek, J. Burdette, J. Kim and S. Xiao; and Start-up Fund from the Environmental and Occupational Health Sciences Institute (EOHSI) at Rutgers University to S. Xiao. This work was also partially supported by The Assistant Secretary of Defense for Health Affairs through the Toxic Exposures Research Program, endorsed by the Department of Defense (DOD) under Award No. HT9425-23-1-0809 (awarded to S. Xiao, NC. Douglas, and Q. Zhang). Opinions, interpretations, conclusions, and recommendations are those of the author and are not necessarily endorsed by the Department of Defense.

