## Supplemental materials for "Ovarian disrupting effects and mechanisms of long- and short-chain per- and polyfluoroalkyl substances in mice"

**Supplemental figures**

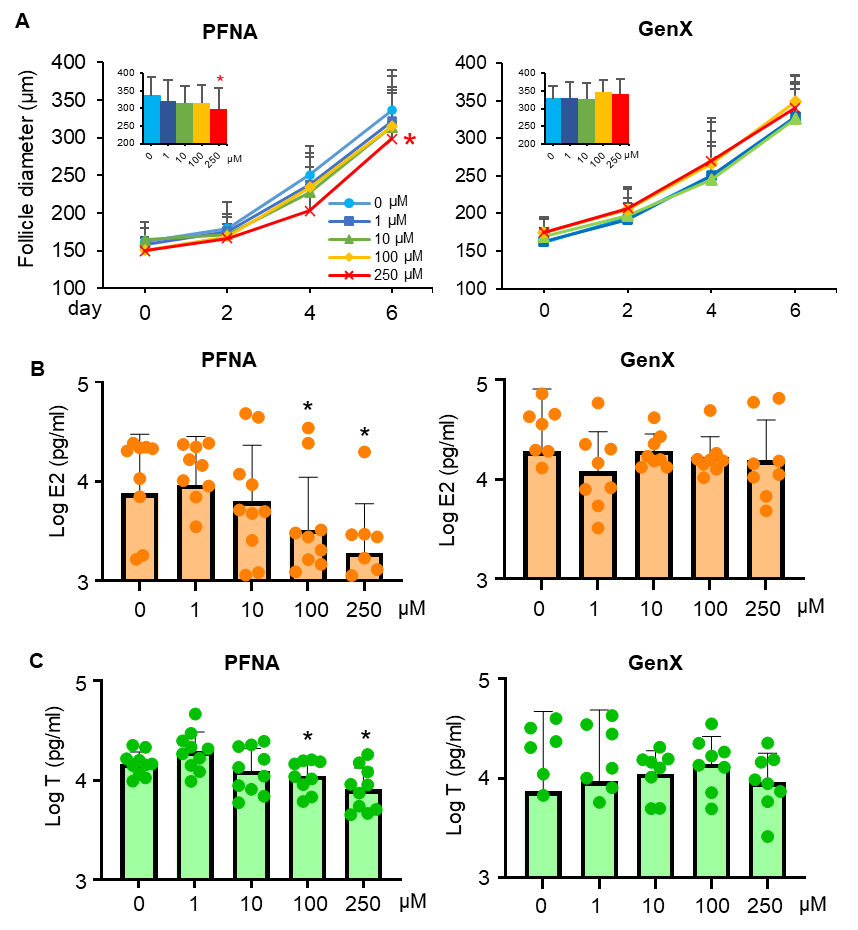

**Supplementary figure S1.** Effects of long-chain and short-chain PFAS on follicle development, steroidogenesis during FSH-dependent maturation window. (A-C) Follicles were treated with PFNA or GenX at range of concentrations from day 2 – day 6. (A) Follicle diameter. (B) Log_10_ estradiol concentration in the conditioned culture media collected from day 6 of eIVFG. (C) Log_10_ testosterone concentration in the conditioned culture media collected from day 6 of eIVFG. Data were analyzed with student’s t-test. N=10 per treatment group. Bars represent standard deviation; **p*˂ 0.05.

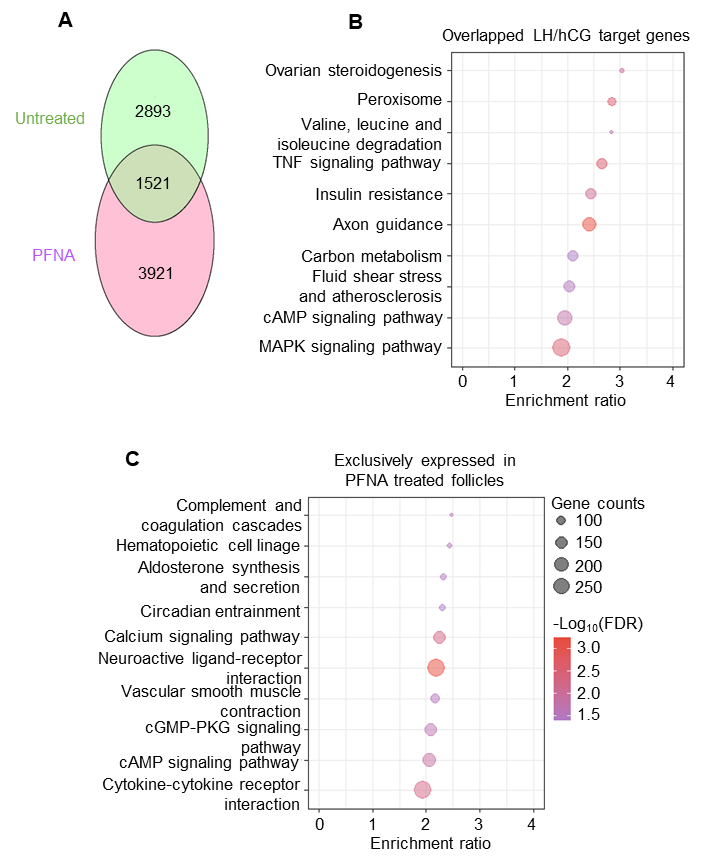

**Supplemental figure S2**. Single follicle RNA-seq data analysis comparing DEGs of untreated follicles to PFNA-treated follicles during ovulation window exposure. (A) Venn diagram of the differentially expressed genes: The number in each circle represents the amount of differentially expressed genes between the different comparisons (untreated vs PFNA-treated). (B-C) KEGG analysis of (B) overlapped LH/hCG target genes, and (C) genes that are exclusively expressed in PFNA-treated follicles. N = 12 for untreated group, N= 10 for PFNA-treated group. Data included in the figure also included in supplemental table 6.

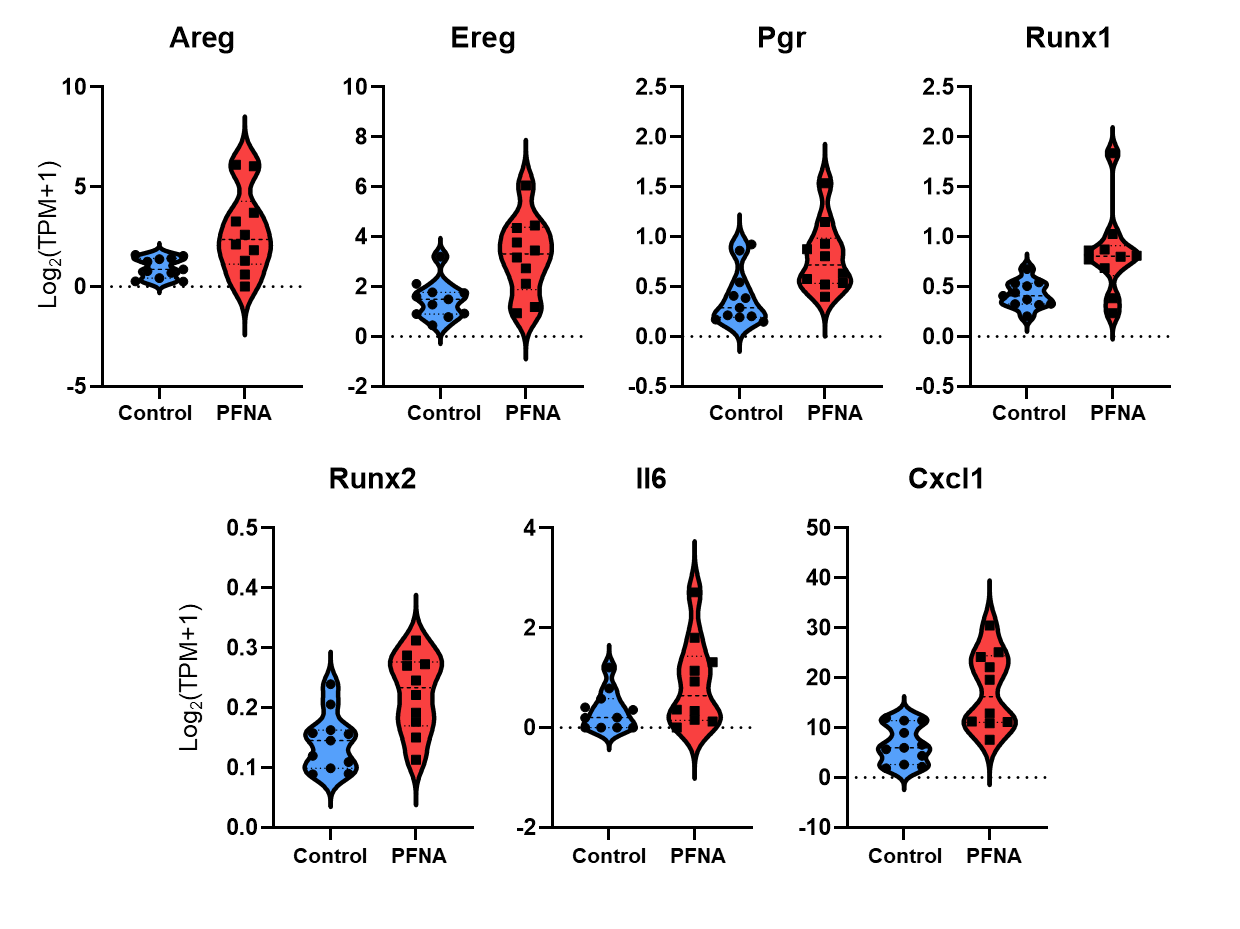

**Supplemental figure S3.** Violin plots of mRNA expression of Pgr-related genes from the Single follicle RNA-seq data analysis (log_2_[TPM+1]) comparing untreated follicles to PFNA-treated follicles (N=8) and control (N=7) during maturation window exposure.

**Supplemental tables**

**Supplemental Table S1.** Primer sequences of examined genes by RT-qPCR

| **Gene** | **Forward Primer (5’-3’)** | **Reverse Primer (5’-3’)** |
| --- | --- | --- |
| *Gapdh* | CATCACTGCCACCCAGAAGACTG | ATGCCAGTGAGCTTCCCGTTCAG |
| *Fshr* | GAGGCAGATGTGTTCTCCAACC | TCGGAGACTGGGAAGATTCTGG |
| *Lhcgr* | GACGCTAATCTCGCTGGAGT | GGCCTGCAATTTGGTGGAAG |
| *Pgr* | TTAAGAGGGCAATGGAAGGGCA | TTTCTCAGACGACATGCTGGG |
| *Ccnd2* | GCAGAAGGACATCCAACCGTA | ACTCCAGCCAAGAAACGGTCCA |
| *PCNA* | CAAGTGGAGAGCTTGGCAATGG | GCAAACGTTAGGTGAACAGGCTC |
| *Pappa* | CAGAAAGCCAGCACCTGTAGCT | GGCAAAGGTCACATGCTGATCC |
| *Inha* | CAGGCTATCCTTTTCCCAGCTAC | AAGTCACCTGGTGGCTGCGTAT |
| *Inhba* | GGAGATAGAGGACGACATTGGC | ACGCTCCACTACTGACAGGTCA |
| *Inhbb* | CTCCGAGATCATCAGCTTTGCAG | GGAGCAGTTTCAGGTACAGCCA |
| *Star* | AAGCTGTGTGCTGGAAGCTC | CTTCCAGTTGAGAACCAAGCAG |
| *Cyp11a1* | TCAAAGCCAGCATCAAGGAGA | TGGCAAAGCTAGCCACCTGTA |
| *Cyp17a1* | CCTGATACGAGGCACTTCTCG | CCAGGACCCAAGTGTGTTCT |
| *Cyp19a1* | CATGGTCCCGGAAACTGTGA | GTAGTAGTTGCAGGCACTTC |
| *Hsd3b1* | AGTGATGGAAAAAGGGCAGGT | GCAAGTTTGTGAGTGGGTTAG |
| *Hsd17b1* | ACTGTGCCAGCAAGTTTGCG | AAGCGGTTCGTGGAGAAGTAG |
| *Ptgs2* | TTCTTTGCCCAGCACTTCAC | CACCTCTCCACCAATGACCTGA |
| *Has2* | GCCGGTCGTCTCAAATTCATC | ACCTCTCACAATGCATCTTGTTC |
| *Tnfaip6* | GCTACAACCCACATGCAAAGG | CTGACCGTACTTGAGCCGAA |
| *Areg* | CACAGCGAGGATGACAAGGA | ATCGTTTCCAAAGGTGCACTG |
| *Ereg* | GCATCCCAGGAGAATCCGAG | GTGTAGCCCACTTCACATCTGC |
| *Btc* | AAACCCACTTCTCTCGGTGC | GCCTTTCTCACAGATGCAGG |
| *Adamts1* | TTGAATGGTGTGAGTGGCGA | CCATCAAACATTCCCCGTGTC |
| *Plat* | AAGAAGCAAGCACTCTCGGG | TCATCTCTGCAGGTCGCTCT |
| *Plau* | CATCCAGTCCTTGCGTGTCT | CCAAGTACACTGCCACCTTCA |
| *Il6* | CCGGAGAGGAGACTTCACAG | TCCACGATTTCCCAGAGA |

**Supplemental Table S2.** Follicle maturation-related genes and their functions

| **Gene** | **Description** | **Functions and references** |
| --- | --- | --- |
| *Fshr* | Follicle stimulating hormone receptor | Activation of FSH target genes and signaling [1, 2] |
| *Lhcgr* | Luteinizing hormone/chorionic gonadotropin receptor | Activation of LH target genes and signaling [3, 4] |
| *Ccnd2* | Cyclin D2 | FSH target genes [5] |
| *PCNA* | proliferating cell nuclear antigen | Cell proliferation related gene; involved in follicle growth [6] |
| *Pappa* | Pregnancy-associated plasma protein A | IGFBP protease; regulation of folliculogenesis and steroidogenesis [7] |
| *Inha* | Inhibin subunit alpha | Synthesis and secretion of ovarian peptide hormones of inhibin and activin and regulation of folliculogenesis [8-11] |
| *Inhba* | Inhibin subunit beta A |  |
| *Inhbb* | Inhibin subunit beta B |  |
| *Star* | Steroidogenic acute regulatory protein | Ovarian steroidogenesis-related gene [12-15] |
| *Cyp11a1* | Cytochrome P450 family 11 subfamily A member 1 |  |
| *Cyp17b1* | Cytochrome P450 family 17 subfamily A member 1 |  |
| *Cyp19a1* | Cytochrome P450 family 19 subfamily A member 1 |  |
| *Hsd3b1* | Hydroxy-delta-5-steroid dehydrogenase, 3 beta- and steroid delta-isomerase 1 |  |
| *Hsd17b1* | Hydroxysteroid 17-beta dehydrogenase 1 |  |

**Supplemental Table S4.** Ovulation-relate genes and their functions

| **Gene** | **Description** | **Ovulatory functions and references** |
| --- | --- | --- |
| *Fshr* | *Follicle stimulating hormone receptor* | Activation of FSH target genes and signaling [1, 2] |
| *Lhcgr* | Luteinizing hormone/chorionic gonadotropin receptor | Activation of LH target genes and signaling [3, 4] |
| *Pgr* | Progesterone receptor | Follicle rupture [16, 17] |
| *Areg* | Amphiregulin | Activation of EGF signaling to regulate cumulus expansion, oocyte maturation, and follicle rupture [18-20] |
| *Ereg* | Epiregulin |  |
| *Btc* | Betacellulin |  |
| *Has2* | Hyaluronan synthase 2 | Cumulus expansion [21] |
| *Tnfaip6* | TNF alpha induced protein 6 | Cumulus expansion [5, 22] |
| *Ptgs2* | Prostaglandin-endoperoxide synthase 2 | Cumulus expansion [23] |
| *Plau* | Plasminogen activator, urokinase | Proteolysis, ECM remodeling, and follicle rupture [24, 25] |
| *Plat* | Plasminogen activator, tissue type | Proteolysis, ECM remodeling, and follicle rupture[16, 24-26] |
| *Adamts1* | A disintegrin and metalloproteinase with thrombospondin motifs 1 | Proteolysis, ECM remodeling, and follicle rupture [16, 26] |
| *Star* | Steroidogenic acute regulatory protein | Ovarian steroidogenesis-related gene [12-15] |
| *Cyp19a1* | Cytochrome P450 family 19 subfamily A member 1 |  |
| *Hsd3b2* | Hydroxy-delta-5-steroid dehydrogenase, 3 beta- and steroid delta-isomerase 2 |  |
| *Cyp17a1* | Cytochrome P450 family 17 subfamily A member 1 |  |
| *Hsd17b1* | Hydroxysteroid 17-beta dehydrogenase 1 |  |
| *Cyp11a1* | Cytochrome P450 family 11 subfamily A member 1 |  |
| *Ccnd2* | Cyclin D2 | FSH target genes [5] |
| *Runx1* | RUNX Family Transcription Factor 1 | Transcription factor [27] |
| *Runx2* | RUNX Family Transcription Factor 2 | Transcription factor [28] |
| *Snap25* | Synaptosome associated protein 25 | PGR target gene [29] |
| *Prkg2* | Protein kinase cGMP-dependent 2 | PGR target gene [30] |
| *Il6* | Interleukin 6 | Proinflammatory factor [31] |
| *Ccl2* | C-C motif chemokine ligand 2 | Proinflammatory factor [32-34] |

**Supplementary table S8:** Characteristics of NHANES reproductive aged women for investigating associations between blood PFOA and PFOS concentrations and long-term amenorrhea.

| **Characteristics** | **Total Sample**  **N (%)** | **Menstruating^1^**  **N (%)** | **Amenorrhea^2^**  **N (%)** | ***p-*value^3^** |
| --- | --- | --- | --- | --- |
| **Total Subjects** | 831 | 799 (95.0) | 32 (5.0) |  |
| **Age, *mean ± SE* (years)** | 31.7 ± 0.36 | 31.6 ±0.35 | 32.9 ±1.60 | 0.5313 |
| **Race/Ethnicity** |  |  |  | 0.0028** |
| Hispanic | 203 (18.1) | 198 (18.6) | 5 (8.7) |  |
| Non-Hispanic White | 297 (58.4) | 278 (54.4) | 19 (80.4) |  |
| Non-Hispanic Black | 186 (12.7) | 180 (12.3) | 6 (8.4) |  |
| Other Race Including Multi-Racial | 145 (10.7) | 143 (10.6) | 2 (2.4) |  |
| **Education Level** |  |  |  | 0.2402 |
| Less than High School | 125 (10.2) | 120 (10.2) | 5 (10.4) |  |
| High School | 155 (18.0) | 146 (17.4) | 9 (29.8) |  |
| More than High School | 551 (71.7) | 533 (72.4) | 18 (59.8) |  |
| **Marital Status** |  |  |  | 0.4061 |
| Married / Living with Partner | 474 (60.1) | 453 (59.5) | 21 (72.4) |  |
| Divorced / Widowed / Separated | 95 (9.9) | 94 (10.2) | 1 (3.7) |  |
| Never Married | 262 (30.0) | 254 (30.3) | 10 (23.8) |  |
| **Covered by Health Insurance** |  |  |  | 0.4728 |
| Yes | 650 (82.5) | 623 (82.8) | 27 (76.2) |  |
| No | 181 (17.5) | 176 (17.2) | 5 (23.8) |  |
| **Poverty Income Ratio, *mean ± SE***  <1  1-1.99  2-2.99  3-3.99  4-4.99  >=5 | 3.57 ± 0.06  224 (19.3)  214 (22.8)  120 (15.1)  92 (12.5)  64 (10.4)  117 (19.9) | 3.38 ± 0.08  229 (20.0)  211 (22.4)  118 (14.9)  88 (11.6)  63 (10.4)  119 (20.6) | 3.90 ± 0.24  5 (9.6)  10 (28.0)  7 (23.8)  5 (22.9)  2 (8.1)  3 (7.6) | 0.2306 |
| **Body Mass Index (kg/m**2)** |  |  |  | n/a |
| Underweight (<18.5) | 21 (2.2) | 21 (2.4) | 0 (.) |  |
| Normal Weight (18.5-24.9) | 268 (33.9) | 256 (34.1) | 12 (31.5) |  |
| Overweight (25.0-29.9) | 193 (23.2) | 185 (22.7) | 8 (32.7) |  |
| Obesity (>30) | 349 (40.6) | 337 (40.8) | 12 (35.9) |  |
| **Ever Smoked** |  |  |  | 0.6980 |
| Yes | 230(30.0) | 222 (30.2) | 8 (25.7) |  |
| No | 601 (70.0) | 577 (69.8) | 24 (74.3) |  |
| **Use hormonal contraception** |  |  |  | 0.5946 |
| Yes | 578 (76.4) | 555 (76.4) | 23 (80.8) |  |
| No | 253 (23.6) | 244 (23.6) | 9 (19.2) |  |

Values for continuous variables are mean +/- the Standard Error of the Mean.

Values for categorical variables are N (unweighted sample counts) and % (weighted sample percentages to account for NHANES survey design).

If the percentages do not equal 100% it is due to rounding.

1. ‘Menstruating’ if answered “Yes” to the question “Have you had at least one menstrual period in the past 12 months? (Please do not include bleedings caused by medical conditions, hormone therapy, or surgeries.)”

2. ‘Long-term amenorrhea’ answered “no” to the question “Have you had at least one menstrual period in the past 12 months? (Please do not include bleedings caused by medical conditions, hormone therapy, or surgeries.)” and answered “Other” or “Don’t know” to the question “What is the reason that you have not had a period in the past 12 months?”

3. *p*-value for categorical variables comes from a chi-squared test, which determines if there is a significant difference between demographics in long-term amenorrhea vs. menstruating women. *p*-values for continuous variables comes from a t-test to determine if there is a significant difference between the means of long-term amenorrhea vs. menstruating. **p*<0.05, ***p*<0.01, ****p*<0.001.
